# Using computational modeling to teach metabolism as a dynamic system improves student performance

**DOI:** 10.1101/2020.02.18.953380

**Authors:** Christine S. Booth, Changsoo Song, Michelle E. Howell, Achilles Rasquinha, Aleš Saska, Resa Helikar, Sharmin M. Sikich, Brian A. Couch, Karin van Dijk, Rebecca L. Roston, Tomáš Helikar

## Abstract

Understanding metabolic function requires knowledge of the dynamics, interdependence, and regulation of biochemical networks. However, current approaches are not optimal to develop the needed mechanistic understanding, and misconceptions about biological processes persist even after graduation. To address these issues, we developed a computational modeling and simulation approach that employs scaffolded learning to teach biochemistry students about the regulation of metabolism. The power of the approach lies in students’ abilities to alter any component or connection in a modeled system and instantly observe the effects of their changes. We find that students who use our approach perform better on biochemistry metabolism questions compared to students in a course that did not use this approach. We also investigated performance by gender and found that our modules may have the potential to increase equity in education. We noted that students are generally positive about the approach and appreciate its benefits. Our modules provide life science instructors with a dynamic and systems-driven approach to teach metabolic regulation and control that improves learning and also equips students with important technical skills.

## Introduction

To ensure that the United States continues to be globally competitive, in science, technology, engineering and math (STEM), students entering the workforce must be adequately prepared to meet emerging challenges. The education community is working to address this need as evidenced by various calls to action developed by national organizations such as the American Association for the Advancement of Science and the National Research Council (1–3). These calls have prompted many educators to re-evaluate the ways in which they approach science education and find ways to identify and develop innovative and evidence-based solutions to educational problems (4–7). The field of biochemistry is no exception, and the American Society for Biochemistry and Molecular Biology (ASBMB) has identified five threshold concepts in biochemistry (8). These threshold concepts are defined as “concepts and skills that, when mastered, represent a transformed understanding of a discipline, without which the learner cannot progress,” and includes the concept of “biochemical pathway dynamics and regulation” (9). Furthermore, ASBMB and others detail the need for improving students’ technical skills. This need, coupled with the inadequacy of traditional methods to teach dynamics and regulation and a shift in life sciences research to incorporate computation, make it increasingly important for life sciences education to equip students with skills to reason mechanistically and quantitatively (10). An important step toward meeting this need has been the broad establishment of computational modeling and simulations as a core competency for undergraduate students (2,3). In this work, we develop and implement a method that increases learning about metabolic pathways as dynamic processes and interconnected networks. Students explore regulation, connectivity, and cellular context by incorporating computational modeling and simulation learning modules in the biochemistry classroom.

Knowledge of metabolism is fundamental to the study of biochemistry. To master metabolism, students have to understand the fundamental concepts, the interrelationship between concepts, and additional “linking ideas” that underlie the interrelationships (11). Schultz (12) highlighted the “learning demand” on students who study metabolic pathways as follows: 1) knowing the particular chemical transformation involved, 2) evaluating the thermodynamics of each step, and 3) comprehending the biological context. The amount of information contained in a single pathway of a metabolic network can quickly overwhelm students, making it difficult to interpret the function and regulation of larger networks or organisms (12–15). Consequently, despite repeated exposure to the same biological system, the difficulties that students face when learning about biochemical pathways ultimately lead to the persistence of misconceptions about metabolism (13,15–18).

A deep understanding of how biological systems function relies on appreciating the dynamic nature of the system. Ultimately, students’ struggles with complex biological processes may be partly attributed to their inability to understand and predict the behavior of systems (i.e. to adopt a systems-thinking perspective). This perspective does not rely on conceptual knowledge and instead requires an analytical approach (19,20). Critical thinking and problem-solving are needed to explain how components interact to support the function of the system. When students adopt a systems-thinking perspective, they must conceptualize systems as interconnected processes, that are themselves nested within larger systems, and whose functions can be mechanistically explained (21,22). Typical instructor practices and student materials may even hamper the adoption of a systems-thinking perspective. For example, most biochemistry textbooks focus on the details of individual enzymatic steps of metabolic pathways and discuss the integrated regulation of metabolic pathways only broadly without explicitly connecting concepts across multiple levels or across multiple chapters.

Computational models of complex biological and biochemical processes and simulations can actively engage students in an experiment-like learning environment (23–25). Computational model-based learning is leveraged in many fields (e.g., chemistry, physics, engineering, biology) and encourages students to make their thinking explicit, which can help students simultaneously hone practical skills, increase content knowledge, and overcome scientific misconceptions (13,23,26–34). Moreover, the need to develop students’ science process skills in general, and modeling abilities specifically, extends beyond the classroom, and the use of models to predict experimental outcomes is a recognized learning goal for biochemistry students (2,3,5).

We hypothesized that teaching metabolism by using computational learning modules with explicit systems-thinking prompts would increase students’ mechanistic understanding of complex biological systems. To target specific learning objectives that were aligned with the ASBMB learning goals, we designed and tested two computational learning modules across a series of upper-level biochemistry courses across two semesters: (1) *Regulation of Cellular Respiration* during Semester 1 and (2) *Regulation of Purine Biosynthesis* during Semester 2 (Tables 1 and 2). During each computational learning module, we used the Predict-Observe-Explain (POE) model of instruction to explicitly ask students to take a systems-thinking perspective (35). We also aligned each learning objective with students’ previously identified difficulties in understanding metabolic processes as systems (Tables 1 and 2). In the first semester, we compared assessment results from each sub-part of the module to those from a course which received typical classroom instruction only (“No module”). Overall, our results indicate that both modules facilitated students’ mechanistic understanding of complex biological systems.

**Table 1.** Alignment of the *Regulation of Cellular Respiration* module learning goals and assessment items with student difficulties, ASBMB learning goals and STH level.

| Topic and associated assessment | Learning objective | Student difficulty (Brown & Schwartz) | Assessment Item | ASBMB learning goal<sup>1</sup> | STH level<sup>2</sup> |
| --- | --- | --- | --- | --- | --- |
| **Glycolysis (Assessment 1.1)** | 1. Mechanistically explain why and how energy charge affects glycolytic intermediates. | Interconnectedness of processes | 1a, 1d, 1g, 1h, 1i | 1,2,3,4 | 5 |
|  | 2. Mechanistically explain the role of glucokinase and hexokinase in glucose absorption. | Scales/nested systems | 1e, 1f, | 1,2,3,4 | 5 |
|  | 3. Contrast mechanisms of regulating glucose absorption to those regulating pyruvate production. | Scales/nested systems | 1b, 1c | 1,2,3 | 6 |
| **TCA (Assessment 1.2)** | 4. Mechanistically explain why and how energy charge regulates TCA cycle intermediates. | Interconnectedness of processes | 2a, 2c, 2e | 1,2,3,4 | 5 |
|  | 5. Mechanistically explain why and how NAD<sup>+</sup>/NADH redox state regulates TCA cycle intermediates. | Interconnectedness of processes | 2d, 2e, 2f | 1,2,3,4 | 5 |
|  | 6. Describe the effect of anaplerotic reactions. | Interconnectedness of processes | 2b, 2g, 2h | 1,2 | 6 |
| **ETC (Assessment 1.3)** | 7. Mechanistically explain the importance of O<sub>2</sub> in cellular respiration. | Scales/nested systems | 3a, 3b, 3j | 1,2,3,4 | 5 |
|  | 8. Mechanistically explain the effect of NAD<sup>+</sup>/NADH redox state on ATP production. | Interconnectedness of processes | 3e, 3f<sup>3</sup>, 3g, 3h | 1,2,3,4 | 5 |
|  | 9. Mechanistically explain the effect energy charge on ATP production. | Interconnectedness of processes | 3c, 3d, | 1,2,3,4 | 5 |
|  | 10. Describe how lactate dehydrogenase maintains glycolysis in the absence of O<sub>2</sub>. | Interconnectedness of processes | 3i | 1,2,3,4 | 6 |

**Table 2.** Alignment of the *Regulation of Purine Biosynthesis* module learning goals and assessment items with student difficulties, ASBMB learning goals and STH level

| Topic and associated assessment | Learning objective | Student difficulty (Brown & Schwartz) | Assessment Item | ASBMB learning goal<sup>1</sup> | STH level<sup>2</sup> |
| --- | --- | --- | --- | --- | --- |
| **Purine biosynthesis (Assessment 2.1)** | 1. Identify and describe individual interactions that contribute to regulation of the *de novo* purine biosynthesis pathway. | Interconnectedness of processes | 1a<sup>3</sup>, 1b, 1c, 1d, 1e<sup>3</sup>, 1f, 1g, 1h, 1i | 3 | 5 |
|  | 2. Mechanistically explain how homeostasis of *de novo* purine biosynthesis is maintained. | Scales/nested systems | 2a, 2b, 2c | 1,2,3,4 | 5 |
|  | 3. Describe how changes in cellular conditions affect the metabolic intermediates of purine biosynthesis. | Scales/nested systems | 3a, 3b, 3c, 3d | 1,2,3,4 | 6 |
|  | 4. Mechanistically explain how mutations in purine biosynthetic enzymes result in metabolic disease. | Scales/nested systems | 4a, 4b, 4c, 4d, 4e, 4f | 1,2,3,4 | 6 |

<sup>1</sup>ASBMB learning goals (8): 1) Relate concentrations of key metabolites to steps of metabolic pathways and describe the roles they play in homeostasis; 2) discuss how chemical processes are compartmentalized in the organism, organ and the cell; 3) summarize the different levels of control (including reaction compartmentalization, gene expression, covalent modification of key enzymes, allosteric regulation of key enzymes, substrate availability and proteolytic cleavage) and relate these different levels of control to homeostasis. 4) Model how perturbations to the steady state can result in changes to the homeostatic state.
<sup>2</sup>Systems Thinking Hierarchy (STH) (41): 1) identify the parts and processes of the system; 2) identify simple relationships; 3) identify dynamic relationships; 4) organize the parts of the system into a framework of relationships; 5) identify whole-system cyclic relationships; 6) recognize hidden dimensions of the system; 7) generalize and problem solve within the system; and 8) think temporally about the system through prediction and retrospection.
<sup>3</sup>Items with negative discrimination were removed from our final analysis

## Results

### Computational learning modules improve student performance on conceptual assessments

To improve students’ abilities to visualize metabolism as a connected network of processes, we used computational model-based learning experiences suitable for students taking an upper-level biochemistry or molecular biology course (Figure 1). The computational learning models and all associated activities (“Module”) were designed using the Cell Collective software and we asked students to predict, observe, and then explain the model’s behavior (36,37). We prompted students to discuss and reflect on their reasoning about the system when they added components or connections to the model.

**Figure 1.**
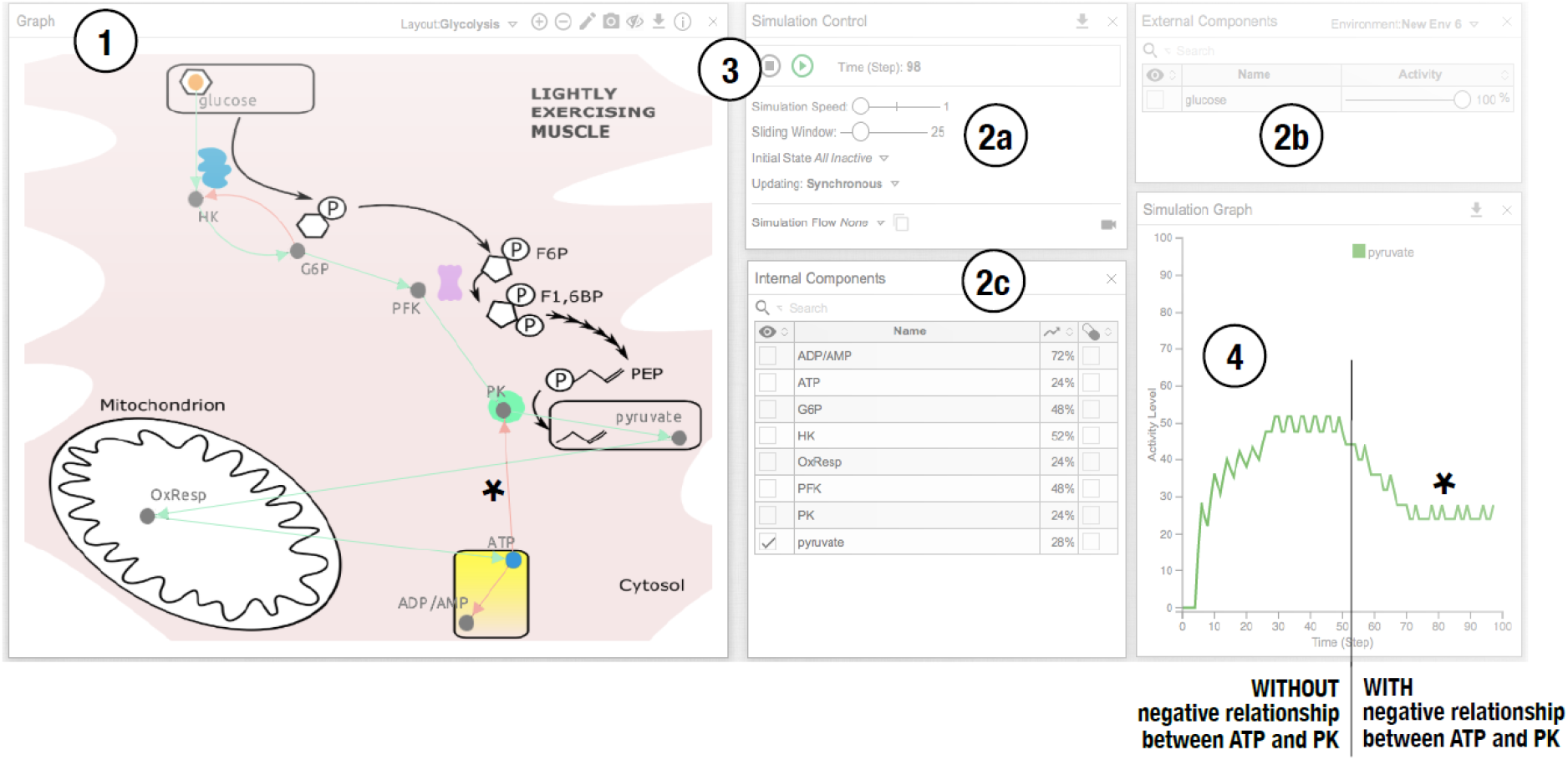
Computational learning modules allow students to take a systems-thinking perspective when using computational models to understand the regulation of metabolic pathways. The Cell Collective web-based software allows students to have an interactive model-based learning experience. Students can (1) edit the computational model by adding components (grey dots) and/or positive or negative relationships (green, red or grey arrows). Students can (2a, b, c) set the simulation parameters, (3) simulate the model’s behavior, and (4) evaluate the effect of changing the model or simulation parameters. For example, to determine the effect of negative allosteric regulation of pyruvate kinase (PK) by ATP, students add a negative relationship between PK and ATP (*) and observe that pyruvate production decreases. In this example, students could also change the level of glucose by adjusting the slider and select additional components to view in the model by checking the box next to them.

We designed and tested two modules in two consecutive courses (Figure 2A and B). The first module, *Regulation of Cellular Respiration*, was integrated in Biochemistry I (Semester 1). It consisted of three sub-parts: *Glycolysis, the Tricarboxylic Acid Cycle (TCA), and the Electron Transport Chain (ETC)*. The second module, *Regulation of Purine Biosynthesis*, was integrated in Biochemistry II (Semester 2). Approximately half of the students in Biochemistry II took the module-integrated Biochemistry I course the previous semester, while the other half were not exposed to the module during Biochemistry I (Figure 2C “Module” vs “No module”).

**Figure 2.**
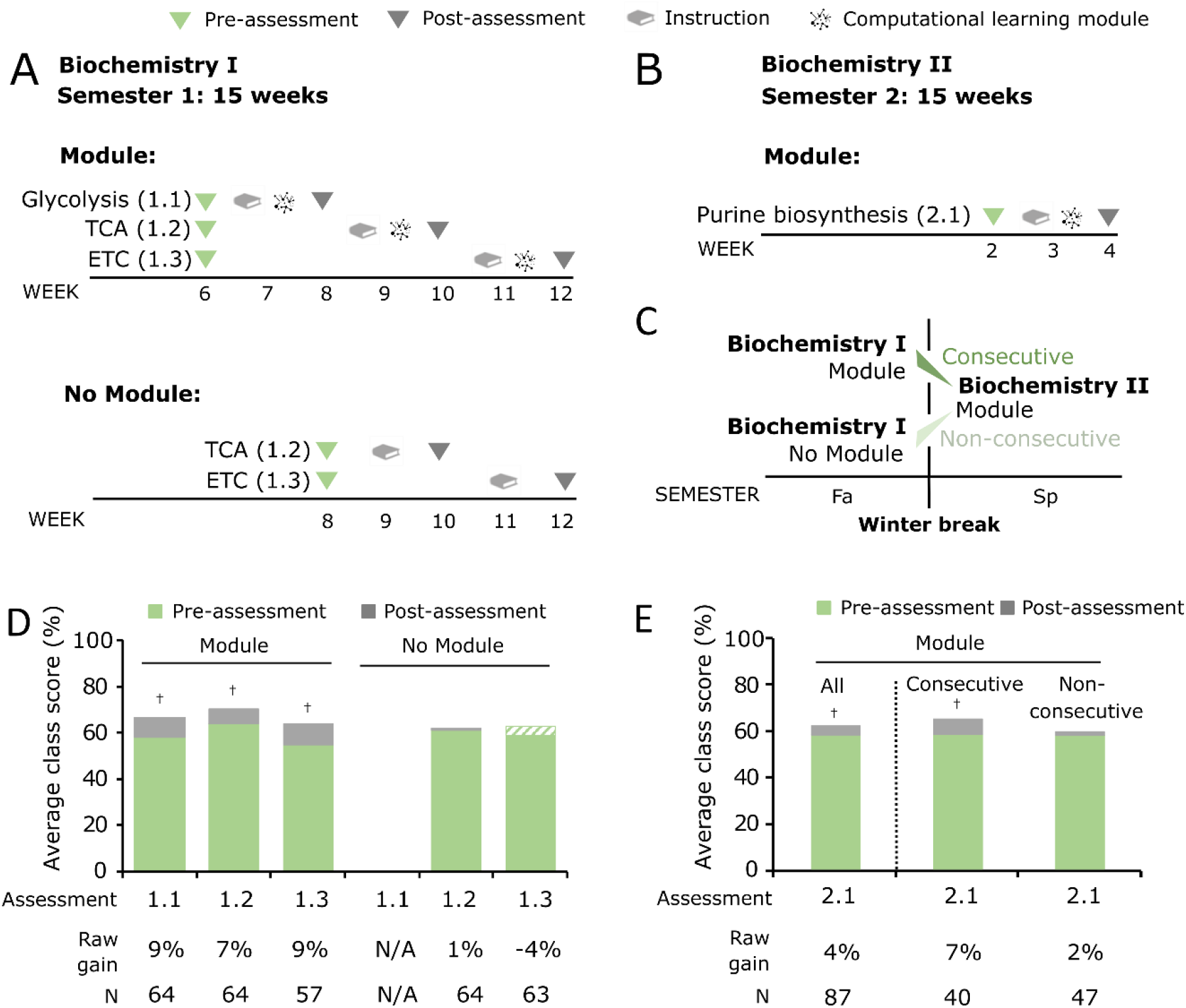
Computational learning modules improve student performance on content assessments for metabolism. A) Diagram of the semester for the “Module” (top) and “No module” (bottom) courses of Biochemistry I. Assessment and instructional timing for *Regulation of Cellular Respiration* is shown. B) Diagram of the semester for the “Module” course of Biochemistry II. Assessment and instructional timing for *Regulation of Purine Biosynthesis* is shown. C) For the purposes of data analysis, students who entered from the Biochemistry I course that used a module were designated to be in the “Consecutive” group (46% of students), while students from the “No module” Biochemistry I course were designated to be in the “Non-consecutive” group (54% of students). D) Course average values of the pre-assessment scores (green) and post-assessment scores (grey) were compared between “Module” and “No module” courses for *Cellular Respiration* (Assessment 1.1: *Glycolysis*, Assessment 1.2: *TCA*, Assessment 1.3: *ETC*). Students in the “No module” course did not complete the *Glycolysis* assessment. Each course was taught by a different instructor. E) Course average values of the pre- and post-assessment scores were compared between “Module” and “No module” courses for *Purine Biosynthesis* (Assessment 2.1). Two-tailed paired t-tests (D: Supporting Table S1 and E: Supporting Table S3) were used to measure significance for pre-versus post-assessment scores: † indicates p<0.05. A green and white striped pattern indicates that the overall post-assessment score was lower than the pre-assessment score.

To determine whether students achieved learning gains after module completion, we analyzed class average scores from pre- to post-assessment. Our data show that integration of the *Regulation of Cellular Respiration* module in Biochemistry I resulted in increased student performance with statistically significant raw learning gains of 9%, 7%, and 9%, for *Glycolysis*, *TCA*, and *ETC* respectively (Figure 2D, Supporting Table S1, “Module”). Conversely, in the “No module” course, gains were statistically indistinguishable at 0% and −4% for *TCA* and *ETC* (Figure 2D, Supporting Table S1, “No module”). To verify the reproducibility of these results, we repeated this experiment in the following academic year (Supporting Figure S1A). Results for the Year 2 replicate were consistent with the first experiment with learning gains of 7%, 6%, and 9% in the “Module” course compared to learning gains of −3%, 0% and 4% in the “No module” course (Supporting Figure S1B, Supporting Table S2).

To determine if this approach of computational module integration could be applied to metabolic pathways that are generally less familiar to students, we integrated a module about the *Regulation of Purine Biosynthesis* in the Biochemistry II course. Student learning gains were similarly measured by evaluating their pre- to post-assessment scores. In addition, we tested whether prior exposure to the Biochemistry I modules impacted student learning gains by taking advantage of the fact that only half of the students in Biochemistry II experienced modules in Biochemistry I. We found that students who were exposed to the modules in Biochemistry I (Figure 2E, “Consecutive”) achieved a significant raw learning gain of 7%, compared to a non-significant gain of 2% for students who were not exposed to modules in Biochemistry I (Figure 2E, “Non-consecutive”). Analysis of pooled Biochemistry II student performance resulted in a 4% learning gain (Figure 2E, “All”, Supporting Table S3).

To account for other factors that could influence these results, we used ANCOVAs that included pre-assessment scores and demographic variables as predictor variables (comparisons of student demographic variables are available in Supporting Tables S4 and S5). When we compared the post-assessment test scores for *Regulation of Cellular Respiration*, we found a significant difference between students in the “Module” and “No module” courses for *TCA* (F(1, 116) = 7.443, p<0.01, partial η^2^ = .060, Supporting Table S6) and *ETC* (F(1, 108) = 7.112, p<0.01, partial η^2^ = .062, Supporting Table S6). When we compared student performance between students who previously completed the module (“Consecutive” group) and students who did not (“Non-consecutive” group) for *Regulation of Purine Biosynthesis* in Biochemistry II, we also found a significant difference in the ANCOVA analysis (F(1, 79) = 8.135, p<0.01, partial η^2^ = .093, Supporting Table S7). Taken together, our results indicate that the modules increase students’ understanding of metabolism. Our results also indicate that previous exposure to modules may support students’ subsequent learning with modules, especially when introducing unfamiliar topics or content.

### Computational learning modules improve student performance on specific learning objectives

To evaluate the effect of the modules on specific learning objectives for *Regulation of Cellular Respiration* in Biochemistry I, we measured the learning gains for each learning objective listed in Table 1 (Figure 3, Supporting Table S8). Similar results were seen in our reproducibility study during Year 2 (Supporting Figure S2, Supporting Table S9). We found that students in the “Module” course often had significant learning gains for those objectives for which the associated concepts were most recently introduced (Figure 3, Table 1). For example, when we introduced the “Energy charge” concept during *Glycolysis*, students achieved a significant learning gain on the associated learning objective, whereas the “Energy charge” concept in the *TCA* and *ETC* assessments did not show the same level of gain. Similarly, when we introduced the “Redox state” concept during *TCA*, students achieved a significant learning gain, yet the same level of gain was not achieved later. The same was true when “Fermentation” was explicitly introduced during the *ETC* portion of the module. This trend was not observed for the “No module” course.

**Figure 3.**
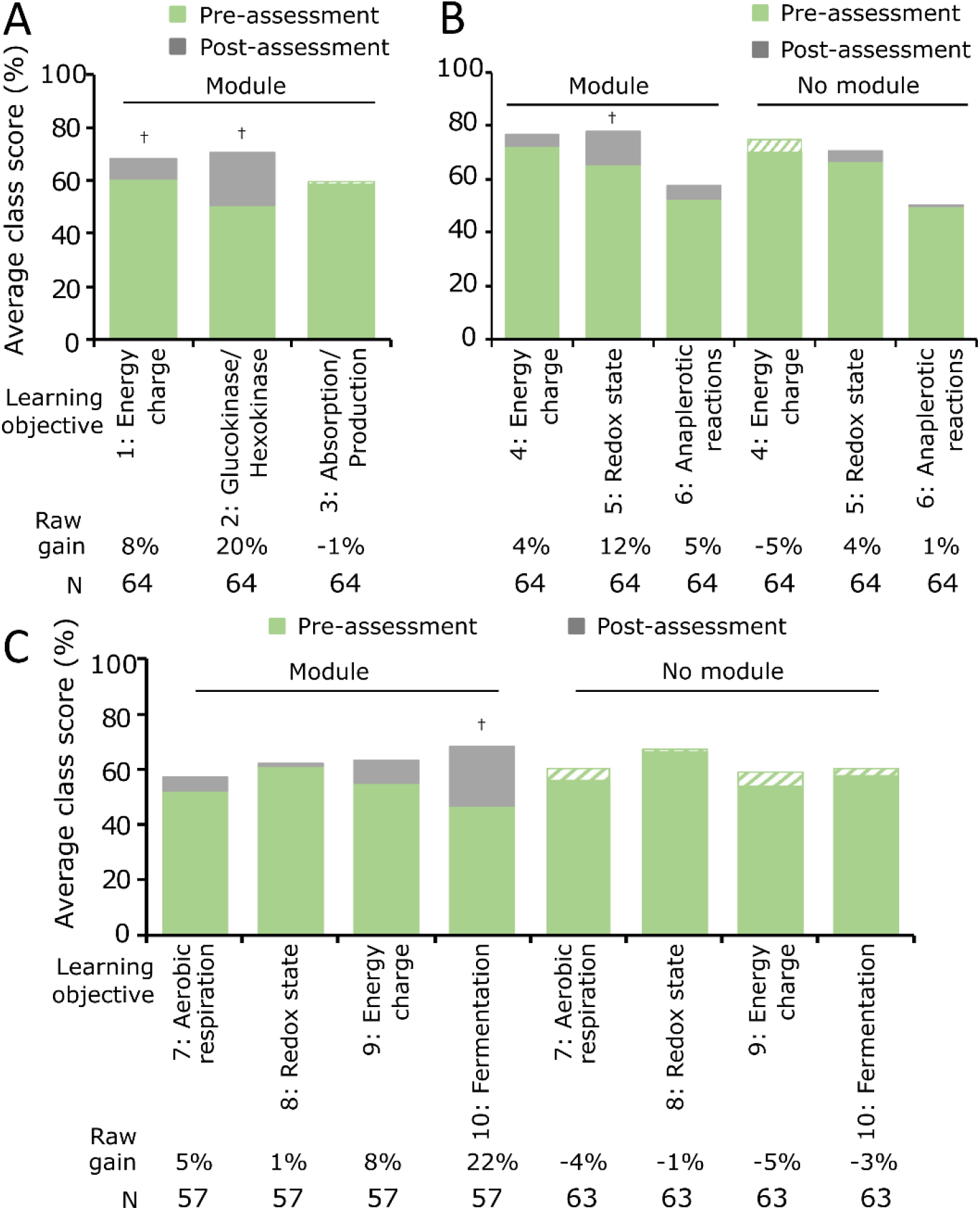
Computational learning modules improve class performance on learning objectives for Cellular Respiration. Average class scores of the pre-assessment scores (green) and post-assessment scores (grey) for each stated learning objective in the *Regulation of Cellular Respiration* module A) Assessment 1.1: *Glycolysis*, B) Assessment 1.2: *TCA*, and C) Assessment 1.3: *ETC* were compared for the “Module” and “No module” courses. Students in the “No module” course did not complete the *Glycolysis* assessment. Each learning objective is numbered, and keywords are provided (refer to Table 1 for detailed objective and corresponding STH level). Two-tailed paired t-tests (Supporting Table S8) were used to measure significance for pre-versus post-assessment scores: † indicates p<0.05

We also analyzed learning gains for each learning objective listed in Table 2 for *Regulation of Purine Biosynthesis* in Biochemistry II (Figure 4, Supporting Table S10). We found that students in the “Consecutive” group achieved significant learning gains on the concept of mutations and disease (the focal concept of the last part of the module); however, students in the “Non-consecutive” group did not achieve significant increases on any learning objectives (Figure 4, Supporting Table S10). Taken together, our results indicate that the modules increase understanding of specific learning objectives, especially when introducing or focusing on important concepts of metabolism.

**Figure 4.**
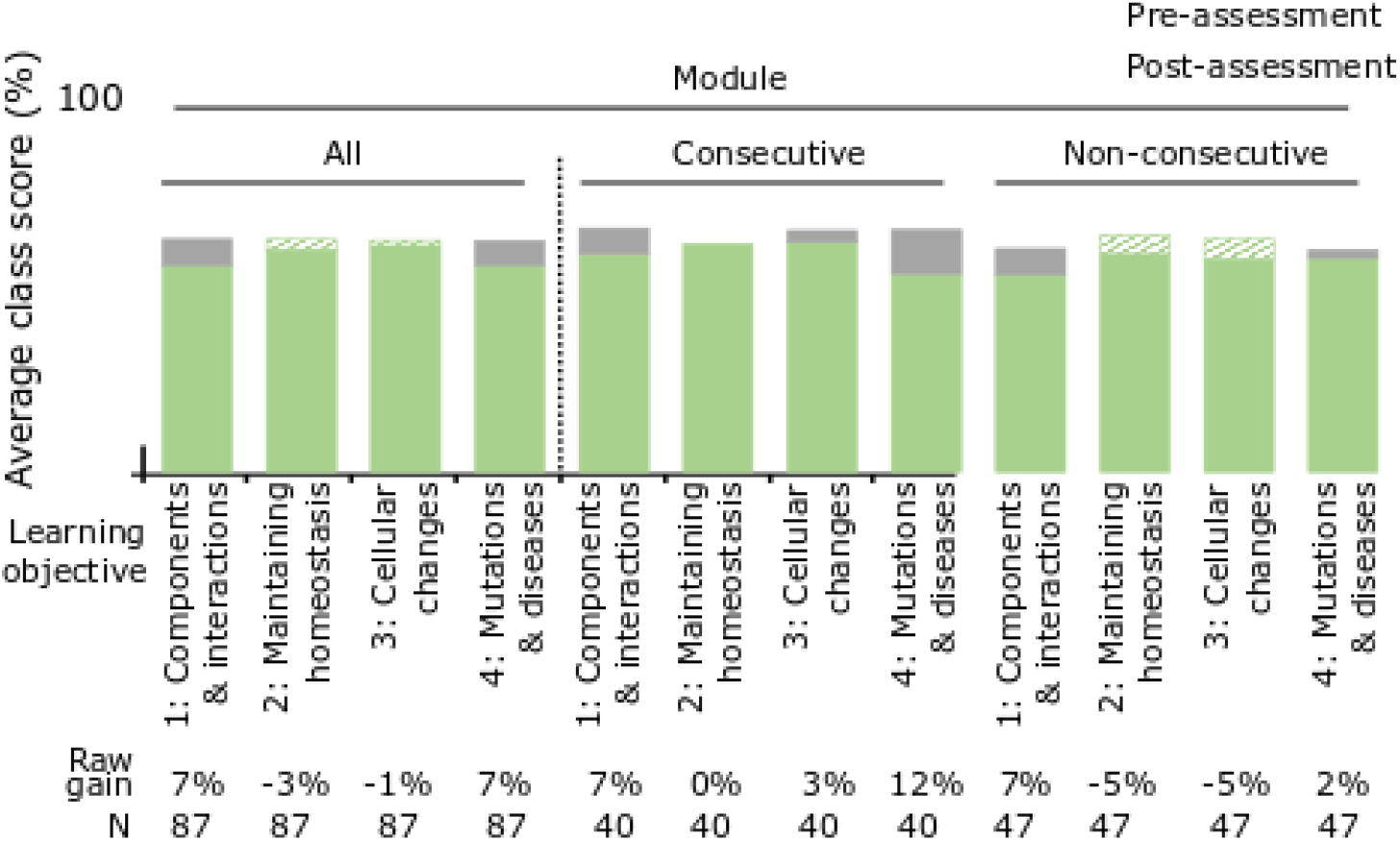
Computational learning modules improve class performance on learning objectives for Purine Biosynthesis. Average class scores of the pre-assessment scores (green) and post-assessment scores (grey) for each stated learning objective in the *Regulation of Purine Biosynthesis* “Module” course (to the left of the dashed line). Assessment results were also compared for the “Consecutive” and “Non-consecutive” groups (to the right of the dashed line). Each learning objective is numbered, and keywords are provided (refer to Table 2 for detailed objective and corresponding STH level). Two-tailed paired t-tests (Supporting Table S9) were used to measure significance for pre- versus post-assessment scores: † indicates p<0.05.

### Repeated computational module interaction may increase learning outcome equity

It has previously been shown that technology use in the classroom can increase gender-based differences in technology-based learning outcomes by impacting students’ attitudes, feelings of inclusion, and learning experiences (38,39). To determine whether there was a difference in the learning gains between male and female students, and to ensure that our modules were not causing a gender gap, we analyzed our results by dividing students in the *Regulation of Cellular Respiration* “Module” and “No module” courses by self-reported gender. We found that male students in the “Module” course achieved a significant pre-post learning gain across the first two module assessments, while female students in the “Module” course only achieved a significant pre-post learning gain for the final assessment in the series (Figure 5A). Notably, neither male nor female students in the “No module” course achieved significant pre-post learning gains (Figure 5B). Interestingly, we noticed a negative trend in learning gains across the semester for male students compared to a positive trend for female students in the “Module” course. This same trend was repeated in the “Module” course for Biochemistry I during our reproducibility study in Year 2 (Supporting Figure S3A). For the “No module” course, we observed that male and female students appeared to trend in the same direction over the course of the semester for both semesters (Figure 5B and Supporting Figure S3B).

**Figure 5.**
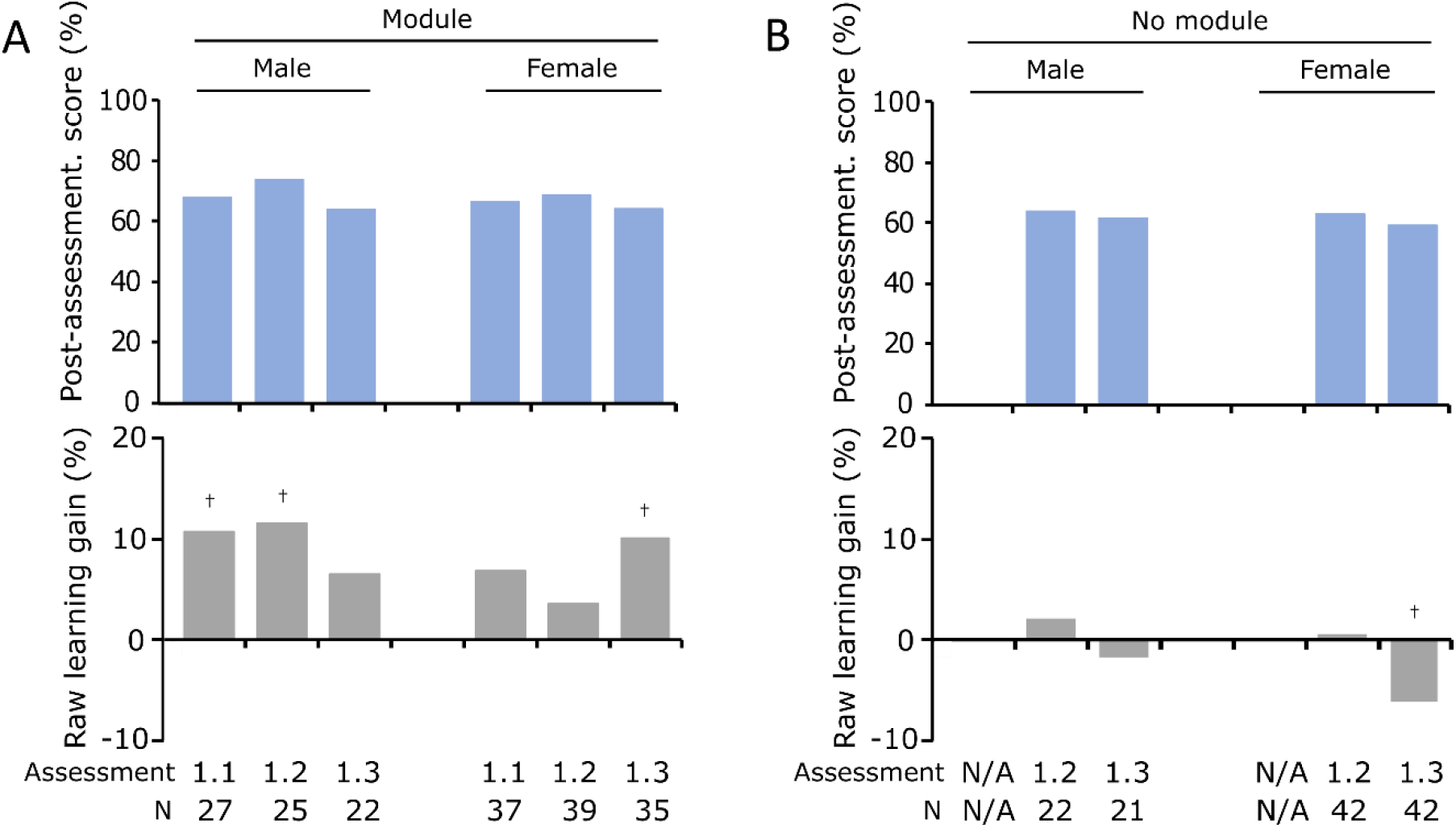
Repeated learning with the computational learning modules may lead to equitable outcomes. Course average values of raw learning gains (grey) for students in the A) Biochemistry I “Module” course, and B) Biochemistry I “No module” course. Learning gains were measured for the topic of Regulation of Cellular Respiration using three assessments (Assessment 1.1 = Glycolysis, Assessment 1.2 = TCA, and Assessment 1.3 = ETC). Two-tailed paired t-tests were used to measure significance for pre- versus post-assessment scores: † indicates p<0.05.

To understand whether the modules differentially impacted performance between the four groups (“Module and male,” “Module and female,” “No module and male,” “No module and female”), we used ANCOVA to compare the test scores after controlling for pre-assessment and other demographic variables. For *TCA*, we detected a significant difference between the groups (F(3,115) = 3.021, p<0.05, partial η^2^ = .073, Supporting Table S11). Specifically, the “Module and male” group was significantly different than the “No module and female” group, indicating that individuals in the “No module and female” group had significantly lower learning gains. For *ETC*, we detected a trend toward a significant difference between the groups (F(3,107) = 2.462, p<0.1, partial η^2^ = .065, Supporting Table S11). Specifically, the “Module and female” group was trending toward a significant difference compared to the “No module and female” group, again indicating that the individuals in the “No module and female” group had significantly lower learning gains. Taken together, the results indicate that females in the “No module” course consistently scored lowest on topics related to metabolism. However, our data also support the idea that, although the modules benefit both male and female students, the benefit to female students increases with repeated exposure.

### Students value the computational learning modules

We used a short survey to determine whether students perceived a learning benefit after completing the modules (Figure 6). In the closed-ended portion of the survey for the *Regulation of Cellular Respiration* module, 54% of students who completed this survey agreed that the module assisted their learning of the material, 45% of the students agreed that they understood what they learned and 48% thought they would remember what they learned (Figure 6A). Sixty percent of students reported that the module reminded them to use a systems-thinking approach that simultaneously considers individual components and the larger system. Likewise, 60% of students reported that the module helped them to understand the effect of feedback loops and environmental conditions, while 58% agreed that the module helped them to understand how the regulation of glycolysis, TCA and ETC are integrated to function as a coherent whole (Figure 6A). The results were similar for Year 2 of Biochemistry I (Supporting Figure S4). For the *Regulation of Purine Biosynthesis* module, we saw a similar, though less dramatic, trend. In addition, students’ perceived learning benefit was comparable between students who were exposed to the module in Biochemistry I and those who were not (Figure 6B and C).

**Figure 6.**
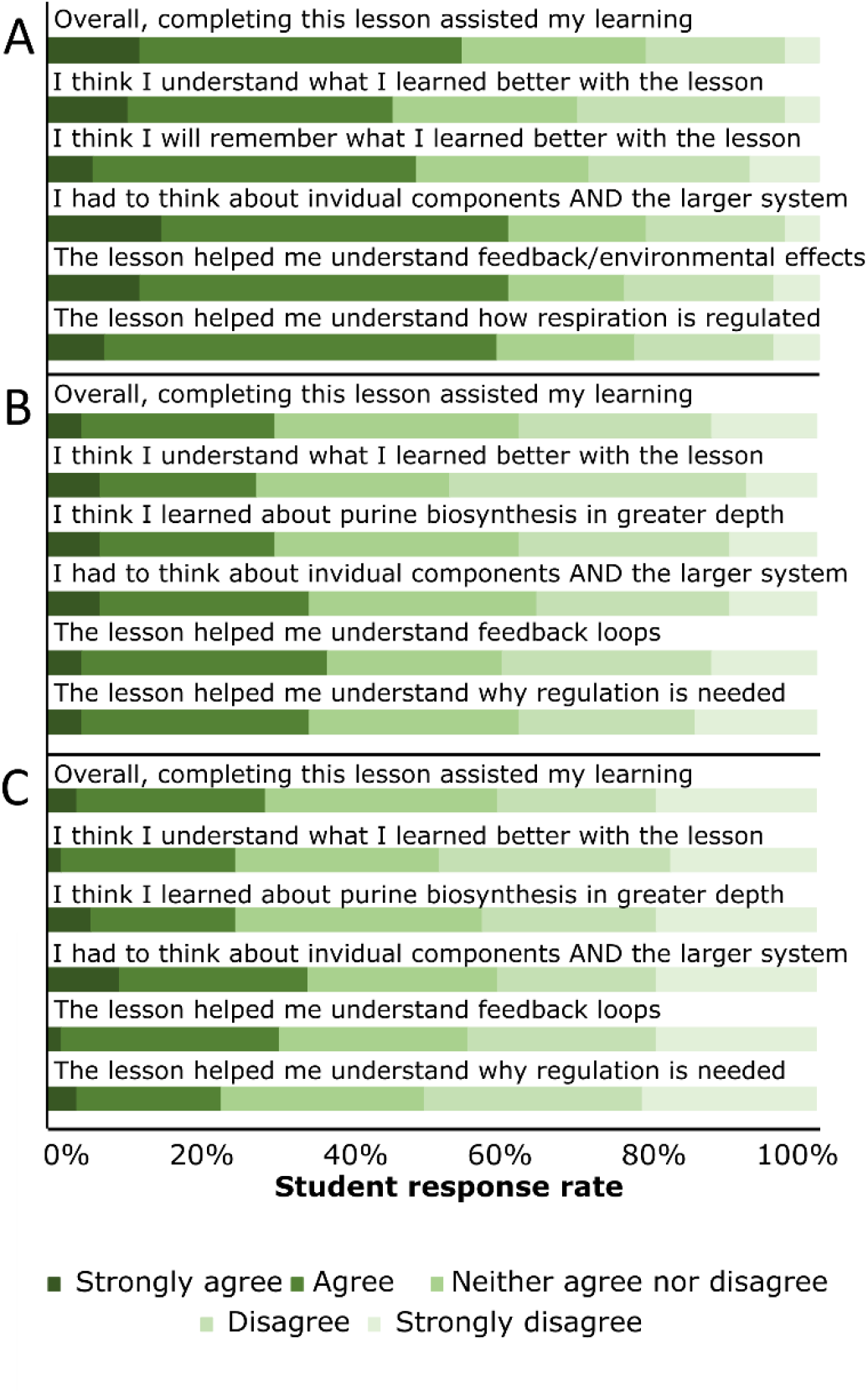
Students valued the modules and report benefits in specific concepts related to systems-thinking. Students in the “Module” courses for A) Biochemistry I *Regulation of Cellular Respiration*, and B) Biochemistry II *Regulation of Purine Biosynthesis* with previous exposure to a module (“Consecutive” group), and C) Biochemistry II *Regulation of Purine Biosynthesis* without previous exposure to a module (“Non-consecutive” group), completed a brief survey about their experiences with the module. Results were reported on a five-point Likert scale.

In the open-ended section of the survey, we asked students to reflect on which aspects of the modules they found to be most beneficial and also the most challenging. Students who completed the *Regulation of Purine Biosynthesis* module had similar responses to the open-ended section of the survey compared to students who completed the *Regulation of Cellular Respiration* module. Benefits included being able to manipulate individual components of the model and directly visualize the effect on the entire system using simulations, seeing the relationships between individual components and multiple processes, and experiencing how the modules foster active learning. One student summarized the importance of simulating the model’s behavior, “The running of the simulations is the most important aspect of the module, at least in my case. It is the only time you are fully able to see what is happening to the levels of different products in the cell and how it affects activity.” Another noted that the module aided their learning by “[seeing how] changing the amount of glucose, LDH, O_2_, and physical exercise alters the production of glycolysis, fermentation, TCA, and ETC and how all of the individual components/metabolites are affected”. A third student noted that the module helped them to “[think] about why the enzymes were connected the way they were and why my predictions were or were not correct”.

Student challenges included keeping track of the number of components and connections involved in the processes as well as feeling concerned about whether the simulations were set up correctly and thus, whether the simulation results were correct. Some students reported frustration about being asked to conceptually evaluate the simulation results. Paradoxically, some students reported positive responses to being asked many conceptual questions about the system components and simulation results.

## Discussion

We responded to national calls for instruction on the dynamics and regulation of biochemical pathways and the use of models to predict experimental outcomes by designing and integrating two interactive computational learning modules in the biochemistry classroom (1–3). We aimed to develop students’ mechanistic understanding of complex biochemical systems and increase their ability to interpret and evaluate them using computational learning modules. We designed the modules to focus on conceptual challenges that undergraduate students face when learning about metabolic pathways.

We found that using the computational learning modules supported students’ learning of concepts and content of metabolism (Figure 2, Supporting Figure S1). Students who learned about the *Regulation of Cellular Respiration* in a “Module” course for Biochemistry I had increased learning gains compared to students who learned about this topic in a “No module” course (Figure 2D, Supporting Figure S1). We also found that students who learned about the unfamiliar topic of *Regulation of Purine Biosynthesis* in a “Module” course for Biochemistry II showed significant learning gains (Figure 2E). Student performance on individual learning objectives revealed that the modules aided students in mastering the mechanistic details of the *Regulation of Cellular Respiration* and *Regulation of Purine Biosynthesis* (Figures 3 and 4, Supporting Figure S2). Our effect sizes derived from statistical models were also consistent with other technology-based learning interventions (24,40). We believe that these results reflect our efforts to make the Cell Collective software and computational learning modules accessible to users of all technical backgrounds through proper user-centric design and scaffolding.

Because our approach hinges on making the systems-thinking perspective explicit, we matched our learning objectives to Assaraf and Orion’s System Thinking Hierarchical (STH) model (41) (Tables 1 and 2). We observed that students achieved significant learning gains more frequently for learning objectives at STH level 5 (5 out of 10 for both modules) compared to objectives as STH level 6 (1 out of 4 for both modules) (Tables 1 and 2, Figures 3 and 4). This could indicate that students may require additional support, exposure, and time investment to achieve understanding at higher STH levels, and instructors may wish to consider adjusting their curriculum to allow as much scaffolding, time and practice as is practical to move student learning to higher STH levels.

We investigated potential differences between male and female participants for two reasons: 1) a large study of thousands of biochemistry students showed that females generally perform worse than males in biochemistry courses (42), and 2) when using technology in the classroom, differences in learning based on gender have been reported (38,39). Consistent with these observations, we found initial differences in learning gains between male and female students learning with the modules (Figure 5A, Supporting Figure 3A). Interestingly, our results indicate that repeated exposures to the modules have the potential to make learning gains more similar for male and female students (Figure 5A, Supporting Figure 3A). Further studies are necessary to identify the specific elements responsible for the observed differences in learning gains trends. To this end, we are designing additional versions of our modules that test each module element separately, and we are expanding our post-module surveys to evaluate the role of students’ confidence and motivation when using technology to learn.

Overall, students self-reported that the modules were valuable learning tools because they 1) reminded them to think about the individual components and the role they play in the larger system, 2) helped them understand the effect of feedback loops and environmental conditions, and 3) helped them appreciate the role of each interaction in the overall regulation of metabolism. During a small focus group conducted by an external evaluator, two students discussed their experience. When asked about the top two most memorable concepts learned during the entire course, one student reported remembering “doing glycolysis, doing the online skills and going through that and learning the up and down regulations...helped me learn how to do the TCA cycle.” When asked how the modules supported student learning, one student noted that “having the [models] as a backup to look at whenever you’re learning such a dense topic is a good way to relearn it besides what’s in the class... It’s a different...hands-on way to look at it, than just having it in front of you and looking at it.” Another student commented on the fact that the systems were so complex that it would be difficult to make predictions about them without first creating a model.

Although students generally valued the computational learning modules, some students were less open to the presented learning approach. These students noted that they would have preferred having lectures or studying the material from the textbook over interacting with the computational models. Our results are consistent with previous findings that classroom interactions and student confidence in the results obtained with models can affect the success of computer model-based instructional approaches (29,34). Some students reported usability issues and commented on their lack of prior knowledge as being challenges to their learning with the modules. To better understand this feedback, we attempted to identify a test group that interacted with an inquiry-based learning environment with high frequency. A small-enrollment course version of biochemistry at a nearby private liberal arts college which emphasizes inquiry-based learning tried the module in the classroom and found similar learning gains. Interestingly, these students rated their learning experience more positively than our students. Our observations are in agreement with findings that students’ curricular exposure shapes their learning profile development, which may determine their readiness for self-directed learning (43). On the usability issues reported, we recognize that technological challenges may be unavoidable with computer-based learning, and we propose that instructors use in-class messaging to encourage students to leave enough time for assistance. Instructors may also increase student buy-in by ensuring close alignment between the modules, class lectures, and exam questions (44,45). Finally, we suggest introducing students to modeling using a familiar system before transitioning to an unfamiliar system, because perceived learning may be lower with unfamiliar systems (Figure 2C-E, Figure 6). Instructors can refer to Supporting File S1 for detailed instructions and incorporation recommendations for a variety of teaching strategies to meet individual course needs.

In summary, students in the life sciences continue to struggle to learn about complex biochemical networks and their regulation (13). At the same time, computational modeling and simulations have emerged as a core component of national education standards in undergraduate life science education (3). The modules described here are provided as a resource for life science instructors to teach metabolic networks using a dynamic, systems-driven approach. Our accompanying assessments also allow instructors to provide formative feedback to their students. Each module asks students to learn by constructing, simulating, and interpreting computational models (37). Students can alter any component or pathway of the process and instantly observe the effect on the modeled system, an approach that closely resembles authentic science practice. Our focus on accessibility has enabled us to make modeling an efficient teaching and learning tool for instructors and students with a broad range of technology or modeling skills. Our modules can be adapted for in-class use, distance-learning, homework and laboratory formats.

## Materials and Methods

### Technology

The modeling and simulation-based learning is facilitated through Cell Collective, a web-based, research-grade software that makes computational modeling accessible to any student and teacher regardless of modeling experience (36,46). Students can alter any network or component of the process and instantly observe the effects of the changes made to the modeled system. In the background, computational models in Cell Collective are mathematically described as probabilistic Boolean control networks (47–49). These models consist of components connected with directed edges. Each component can represent a variety of elements ranging from a single enzyme or metabolite to an entire process, depending on the scope of the model and the level of abstraction. The directed edges correspond to biochemical interactions (direct or indirect) among the components (e.g., Glucose-6-phosphate activates Glucose-6-phosphate dehydrogenase IF/WHEN NAD+ is active) (50). The activity of a component is determined by its regulatory mechanism, reflecting the activity of other directly interacting components. The components’ simulation output provides a semi-quantitative measure to describe the relative level of activity, rather than a specific biological measure such as abundance or concentration. This allows students to observe the effects of their changes to the model. Cell Collective can be accessed directly via a web-browser (i.e., no installation is needed) by visiting https://cellcollective.org and selecting “Learning - Get started” on the home page. Users can access all content without registration and users who create free accounts can save their work.

### Module design

To encourage students to take a systems perspective and develop their mechanistic understanding of complex biological systems, we designed computational learning modules suitable for upper-level biochemistry or molecular biology courses. The first module, *Regulation of Cellular Respiration*, consisted of three sub-parts: *Glycolysis* (Assessment 1.1), *TCA* (Assessment 1.2), and *ETC* (Assessment 1.3), and addressed 1) how the energy charge- and redox-status of the cell regulate glycolysis and the TCA cycle, 2) how the ETC is integrated into this system, and 3) how the system maintains homeostasis despite changes to the environment (e.g. oxygen availability) (Supporting Files S2-S4). The second module, *Regulation of Purine Biosynthesis* (Assessment 2.1) addressed 1) how the regulatory mechanisms of the purine biosynthesis pathway allows the cell to maintain homeostasis despite changes to the environment, and 2) how mutations disrupt the cell’s ability to maintain homeostasis (Supporting Files S5-S7).

Instructional scientists and researchers typically emphasize the benefits of having students build, evaluate, and revise their own models as opposed to simply using expert constructed models (30,51). However, due to the complexity of the systems under study, and the time that would be required to fully model and troubleshoot the behavior of the systems, we elected to use an intermediate “model elaboration” approach, in which we provided students with key components, asked them to add in known regulatory relationships, and reason through the effects of these individual relationships on the function of the entire system (32). Using the learning objectives as a guide, we developed computational models up to three months before the planned class to allow sufficient time to adjust the module and optimize the models to fit the module design. We used textbook sources to identify critical components to be included in each computational model, and manually curated published evidence for regulation. When available, we also used published literature to confirm model outputs. The fully annotated models including literature references are available in the Cell Collective software.

Incorporated into each model, we created a series of interactive activities that provide students with informational prompts, instructions, and questions as they interact with the model building and simulation components of the software (Figure 1). For example, in *Regulation of Cellular Respiration: Glycolysis*, students are provided with a partially built model that is missing important allosteric feedback relationships (Figure 1-1). Students can edit the model, evaluate and predict what effect their edits will have, and simulate the model’s behavior to test whether their predictions were accurate (Figure 1-2 to 1-4).

We used an iterative approach to test and refine the module activities and assessment. For each module, we conducted a think-aloud exercise with one to four senior biochemistry or graduate students focused on usability testing. During the sessions we noted what participants were saying and doing to ensure that we were achieving the desired interaction with the module. When it was obvious that participants were struggling, we engaged them directly to understand the source of their difficulties. This process helped us to develop activities that could be used as stand-alone assignments to reduce instructor burden and increase benefits for distance learning students.

### Implementation

For each module, we followed the same general format of 1) pre-assessment, 2) instruction and module activities, 3) post-assessment (Supporting Files S2-S7). To prepare for class, the instructor and teaching assistants completed the module. Approximately a week before the simulation module was started in-class, students individually completed the closed-ended online pre-assessment (Figures 2A and B, Supporting Files S4 and S7). For our assessments, we used multiple-true-false (MTF) questions that consist of a question stem that is presented together with a series of statements that are evaluated by students as being true or false. We selected MTF questions because they can reveal student misconceptions that remain undetected in free-response and multiple-choice question formats (53–55). The number of statements for each question is listed in Tables 1 and 2.

Before class, students were asked to complete a Cell Collective training module to familiarize themselves with the technology and modeling concepts and to complete the pre-assessment questions online. Students were introduced to computational modeling as it relates to the topic through a mini-lecture. Students worked in groups of two to four, and we used whole-class clicker questions and peer instruction to ensure that students were on target with major concepts and to identify and resolve student misunderstandings and technology issues (52). We required students to complete any unfinished activities as homework. In the case of *Regulation of Cellular Respiration*, students only started the *Glycolysis* part of the module during class and completed all remaining activities as homework (i.e., *TCA* and *ETC* were completed entirely as homework assignments over the course of six weeks). Introduction of the modules did not appreciably alter the instructional schedule, and diagrams of the timing of module activities and assessments during the course of the semester are shown (Figures 2A and B).

After completing the modules as homework, students answered the same post-assessment questions online to evaluate their learning gains. Students also completed a short survey about their experiences with the models (Supporting Files S4, S7-S9).

The Supporting Information contains additional materials and information necessary to implement the modules, including slides for mini-lectures (Supporting Files S2 and S5), instructor guides (Supporting Files S3 and S6), assessment questions (Supporting Files S4 and S7), and student experience survey questions (Supporting Files S8 and S9).

### Data Collection, Participants and Data Analysis

We implemented the computational learning modules in two large-enrollment senior-level undergraduate biochemistry courses. The aforementioned courses comprise a two-part series (here called Biochemistry I and Biochemistry II) that are typically taken in sequence. Both classes are required for biochemistry majors and contain a large pre-health population at a research-intensive university.

The *Regulation of Cellular Respiration* module was implemented in Biochemistry I (N = 107) and the *Regulation of Purine Biosynthesis* module was implemented in Biochemistry II (N = 142). Here, we report only the results using data from consenting students for whom we had demographic information and who completed both the pre- and post-assessments. For *Regulation of Cellular Respiration*, N = 64, 64, 57 for Glycolysis, *TCA*, and *ETC* assessments and for *Regulation of Purine Biosynthesis*, N = 87. Each component of the module (pre-assessment, module activities, and post-assessment) was graded based on completion in both “Module” courses. For the *Regulation of Cellular Respiration*, we compared the learning gains from each class to those of students in another section of the same course (Biochemistry I) taught by a different instructor at the same university that did not complete the modules (the “No module” course). The “No module” course was taught in a large-enrollment class during the same semester as the “Module” course. Logistically, the “No module” course was unable to complete an equivalent *Glycolysis* assessment and took the pre-assessment slightly later in the semester compared to the module course. For the “No module” course, N = 64 and 63 for the *TCA* and *ETC* assessments.

Approximately equal numbers of students from the “Module” and “No module” courses for Biochemistry I were subsequently exposed to modules in Biochemistry II. Students who were exposed to the module in Biochemistry I were designated as “Consecutive” students, while students who were not exposed to the module in Biochemistry I were designated as “Non-consecutive” students (Figure 2C).

Supporting Table S4 contains participant’s demographic profiles for the “Module” and “No module” courses in Biochemistry I. We noted a significant difference (*p* < 0.01) between the cumulative GPA in the “Module” course (3.74, SD = 0.28) and “No module” course (3.41, SD = 0.85). Supporting Table S5 contains participant’s demographic profiles for the “Consecutive” and “Non-consecutive” groups in Biochemistry II. Again, we noted a significant difference (*p* < 0.05) between the cumulative GPA in the “Consecutive” (3.64, SD = 0.39) and “Non-consecutive” groups (3.81, SD = 0.18).

We used item analysis to evaluate the quality of our assessments. We combined student responses from the “Module” and “No module” courses to determine item difficulty and discrimination for the pre- and post-assessment scores. We excluded items with negative discrimination on the post-assessment score from our analysis and removed them from our future assessments.

For data analysis we first calculated the raw learning gain (post-assessment score – pre-assessment score) for each student. Then, for each assessment, we calculated the mean raw learning gain for the entire class. For all courses (“Module” and “No module”), we determined whether students significantly improved from pre- to post-assessment by performing a two-tailed paired t-tests on individual student performance. We used a similar approach to analyze student learning gains for each learning objective.

Next, we used IBM SPSS 23.0 to conduct a one-way ANCOVA to determine whether a statistically significant difference existed on the post-assessment score for each assessment between the “Module” and “No module” courses in Biochemistry I when controlling for the covariates (pre-test score and cumulative GPA) and the demographic variables (i.e., gender, native English speaker, parents’ college education, and the extent of education self-funding). Post hoc analyses for pairwise comparisons were performed with a Bonferroni adjustment. We followed the same approach to evaluate the significance of differences between “Consecutive” and “Non-consecutive” groups in Biochemistry II. When analyzing differences between males and females, we followed the same approach after also dividing the groups based on self-reported gender (“Module and male”, “Module and female”, “No module and male”, and “No module and female”).

## Acknowledgments

We thank Audrey Crowther, Thuan Long, Jon Dietz, Daisy Guiza Beltran, James Orf, Grant Ozaki, Brady Caverzagie, Ben Plambeck, Olivia Makos, and Dr. Lisa Briona for feedback on the modules. We thank Drs. Edward N. Harris, and Xinghui Sun for administering assessments in their courses. This work was supported by the National Science Foundation grant number NSF DUE-1625804 and was classified as exempt from IRB review.

## References

1. National Research Council, “A framework for K-12 science education: Practices, crosscutting concepts, and core ideas” in (The National Academies Press, 2012).

2. National Research Council, “Next generation science standards: For states, by states” in (The National Academies Press, 2013).

3. American Association for the Advancement of Science, “Vision and change in undergraduate biology education: a call to action” (2011).

4. M. Kramer, D. Olson, J. D. Walker, Design and Assessment of Online, Interactive Tutorials That Teach Science Process Skills. CBE—Life Sciences Education 17, ar19–ar19 (2018).

5. H. B. White, M. A. Benore, T. F. Sumter, B. D. Caldwell, E. Bell, What skills should students of undergraduate biochemistry and molecular biology programs have upon graduation? Biochemistry and molecular biology education 41, 297–301 (2013).

6. M. E. Howell, et al., Visualizing the Invisible: A Guide to Designing, Printing, and Incorporating Dynamic 3D Molecular Models to Teach Structure–Function Relationships. J Microbiol Biol Educ 19 (2018).

7. M. E. Howell, et al., Student Understanding of DNA Structure–Function Relationships Improves from Using 3D Learning Modules with Dynamic 3D Printed Models. Biochemistry and Molecular Biology Education 47, 303–317 (2019).

8. J. T. Tansey, et al., Foundational concepts and underlying theories for majors in biochemistry and molecular biology. Biochemistry and Molecular Biology Education 41, 289–296 (2013).

9. J. Loertscher, D. Green, J. E. Lewis, S. Lin, V. Minderhout, Identification of threshold concepts for biochemistry. CBE—Life Sciences Education 13, 516–528 (2014).

10. National Science and Technology Council Committee on STEM Education, “Charting a Course for Success: America’s Strategy for STEM Education” (2018).

11. T. Waheed, A. M. Lucas, Understanding interrelated topics: photosynthesis at age photosynthesis at age 14+. Journal of Biological Education 26, 193–199 (1992).

12. E. Schultz, A guided discovery approach for learning metabolic pathways. Biochemistry and Molecular Biology Education 33, 1–7 (2005).

13. M. H. Brown, R. S. Schwartz, Connecting photosynthesis and cellular respiration: Preservice teachers’ conceptions. Journal of Research in Science Teaching: The Official Journal of the National Association for Research in Science Teaching 46, 791–812 (2009).

14. C. E. Hmelo-Silver, M. G. Pfeffer, Comparing expert and novice understanding of a complex system from the perspective of structures, behaviors, and functions. Cognitive Science 28, 127–138 (2004).

15. C. W. Anderson, T. H. Sheldon, J. Dubay, The effects of instruction on college nonmajors’ conceptions of respiration and photosynthesis. Journal of Research in Science teaching 27, 761–776 (1990).

16. C. D. Wilson, et al., Assessing students’ ability to trace matter in dynamic systems in cell biology. CBE—Life Sciences Education 5, 323–331 (2006).

17. J. A. Michael, et al., Undergraduate students’ misconceptions about respiratory physiology. Advances in Physiology Education 277, S127 (1999).

18. J. M. Dauer, J. H. Doherty, A. L. Freed, C. W. Anderson, Connections between Student Explanations and Arguments from Evidence about Plant Growth. Cell Biology Education 13, 397–409 (2014).

19. R. D. Arnold, J. P. Wade, A Definition of Systems Thinking: A Systems Approach. Procedia Computer Science 44, 669–678 (2015).

20. R. P. Verhoeff, M.-C. P. J. Knippels, M. G. R. Gilissen, K. T. Boersma, The theoretical nature of systems thinking. Perspectives on systems thinking in biology education. Frontiers in Education 3, 1–11 (2018).

21. E. Abrams, S. Southerland, The how’s and why’s of biological change: How learners neglect physical mechanisms in their search for meaning. International Journal of Science Education 23, 1271–1281 (2001).

22. L. K. Wright, J. N. Fisk, D. L. Newman, DNA→ RNA: What do students think the arrow means? CBE—Life Sciences Education 13, 338–348 (2014).

23. W. Riess, C. Mischo, Promoting systems thinking through biology lessons. International Journal of Science Education 32, 705–725 (2010).

24. S. Bayraktar, A meta-analysis of the effectiveness of computer-assisted instruction in science education. Journal of research on technology in education 34, 173–188 (2001).

25. N. Rutten, J. T. van der Veen, W. R. van Joolingen, Inquiry-Based Whole-Class Teaching with Computer Simulations in Physics. International Journal of Science Education 37, 1225–1245 (2015).

26. J. Tripto, O. B.-Z. Assaraf, M. Amit, Mapping What They Know: Concept Maps as an Effective Tool for Assessing Students’ Systems Thinking. American Journal of Operations Research 03, 245–258 (2013).

27. J. Stewart, J. L. Cartier, C. M. Passmore, Developing understanding through model-based inquiry. How students learn, 515–565 (2005).

28. S. D. Hester, et al., Authentic Inquiry through Modeling in Biology (AIM-Bio): An Introductory Laboratory Curriculum That Increases Undergraduates’ Scientific Agency and Skills. CBE— Life Sciences Education 17, ar63–ar63 (2018).

29. L. L. Liang, G. W. Fulmer, D. M. Majerich, R. Clevenstine, R. Howanski, The effects of a model-based physics curriculum program with a physics first approach: A causal-comparative study. Journal of Science Education and Technology 21, 114–124 (2012).

30. G. King, H. E. Bergan-Roller, N. J. Galt, T. Helikar, J. T. Dauer, Modelling activities integrating construction and simulation supported explanatory and evaluative reasoning. International Journal of Science Education, 1764–1786 (2019).

31. H. E. Bergan-Roller, N. J. Galt, J. T. Dauer, T. Helikar, Discovering Cellular Respiration with Computational Modeling and Simulations. CourseSource (2017).

32. H. E. Bergan-Roller, N. J. Galt, C. J. Chizinski, T. Helikar, J. T. Dauer, Simulated Computational Model Lesson Improves Foundational Systems Thinking Skills and Conceptual Knowledge in Biology Students. BioScience 68, 612–621 (2018).

33. G. Martinez, F. L. Naranjo, A. L. Perez, M. I. Suero, P. J. Pardo, Comparative study of the effectiveness of three learning environments: Hyper-realistic virtual simulations, traditional schematic simulations and traditional laboratory. Physical Review Special Topics-Physics Education Research 7, 20111–20111 (2011).

34. S. J. Streicher, K. West, D. M. Fraser, J. M. Case, C. Linder, Learning through simulation: Student engagement. Chemical Engineering Education 39, 288–295 (2005).

35. M. Kearney, D. F. Treagust, S. Yeo, M. G. Zadnik, Student and teacher perceptions of the use of multimedia supported predict–observe–explain tasks to probe understanding. Research in Science Education 31, 589–615 (2001).

36. T. Helikar, et al., The Cell Collective: Toward an open and collaborative approach to systems biology. BMC Systems Biology 6, 96–96 (2012).

37. T. Helikar, et al., Integrating Interactive Computational Modeling in Biology Curricula. PLOS Computational Biology 11, e1004131–e1004131 (2015).

38. B. J. Young, Gender differences in student attitudes toward computers. Journal of research on computing in education 33, 204–216 (2000).

39. I. Heemskerk, G. ten Dam, M. Volman, W. Admiraal, Gender inclusiveness in educational technology and learning experiences of girls and boys. Journal of Research on Technology in Education 41, 253–276 (2009).

40. D. A. Cook, et al., Comparative effectiveness of instructional design features in simulation-based education: systematic review and meta-analysis. Medical teacher 35, e867–98 (2013).

41. O. B.-Z. Assaraf, N. Orion, Development of system thinking skills in the context of earth system education. Journal of Research in Science Teaching 42, 518–560 (2005).

42. M. M. Rauschenberger, R. D. Sweeder, Gender performance differences in biochemistry. Biochemistry and Molecular Biology Education 38, 380–384 (2010).

43. C. Kell, R. Van Deursen, Student learning preferences reflect curricular change. Medical teacher 24, 32–40 (2002).

44. K. R. Brazeal, T. L. Brown, B. A. Couch, Characterizing student perceptions of and buy-in toward common formative assessment techniques. CBE—Life Sciences Education 15, ar73–ar73 (2016).

45. G. Wiggins, G. P. Wiggins, J. McTighe, Understanding by Design (ASCD, 2005).

46. T. Helikar, B. Kowal, J. A. Rogers, A cell simulator platform: the cell collective. Clinical Pharmacology & Therapeutics 93, 393–395 (2013).

47. M. M, C. F, K. D, N. K, D. S, Cell cycle-regulated transcription of the CLB2 gene is dependent on Mcm1 and a ternary complex factor. Mol Cell Biol 15, 3129–3137 (1995).

48. W. Abou-Jaoudé, et al., Logical modeling and dynamical analysis of cellular networks. Frontiers in genetics 7, 94–94 (2016).

49. T. Helikar, J. Konvalina, J. Heidel, J. A. Rogers, Emergent decision-making in biological signal transduction networks. Proceedings of the National Academy of Sciences 105, 1913–1918 (2008).

50. T. Helikar, et al., Bio-Logic Builder: A Non-Technical Tool for Building Dynamical, Qualitative Models. PLOS ONE 7, e46417 (2012).

51. J. D. Gobert, B. C. Buckley, Introduction to model-based teaching and learning in science education. International Journal of Science Education 22, 891–894 (2000).

52. B. A. Couch, J. K. Hubbard, C. E. Brassil, Multiple–true–false questions reveal the limits of the multiple–choice format for detecting students with incomplete understandings. BioScience 68, 455–463 (2018).

53. C. E. Brassil, B. A. Couch, Multiple-true-false questions reveal more thoroughly the complexity of student thinking than multiple-choice questions: a Bayesian item response model comparison. IJ STEM Ed 6, 16 (2019).

54. J. K. Hubbard, M. A. Potts, B. A. Couch, How Question Types Reveal Student Thinking: An Experimental Comparison of Multiple-True-False and Free-Response Formats. LSE 16, ar26 (2017).

55. C. H. Crouch, E. Mazur, Peer instruction: Ten years of experience and results. American journal of physics 69, 970–977 (2001).

