## Supporting Information for "Using computational modeling to teach metabolism as a dynamic system improves student performance"

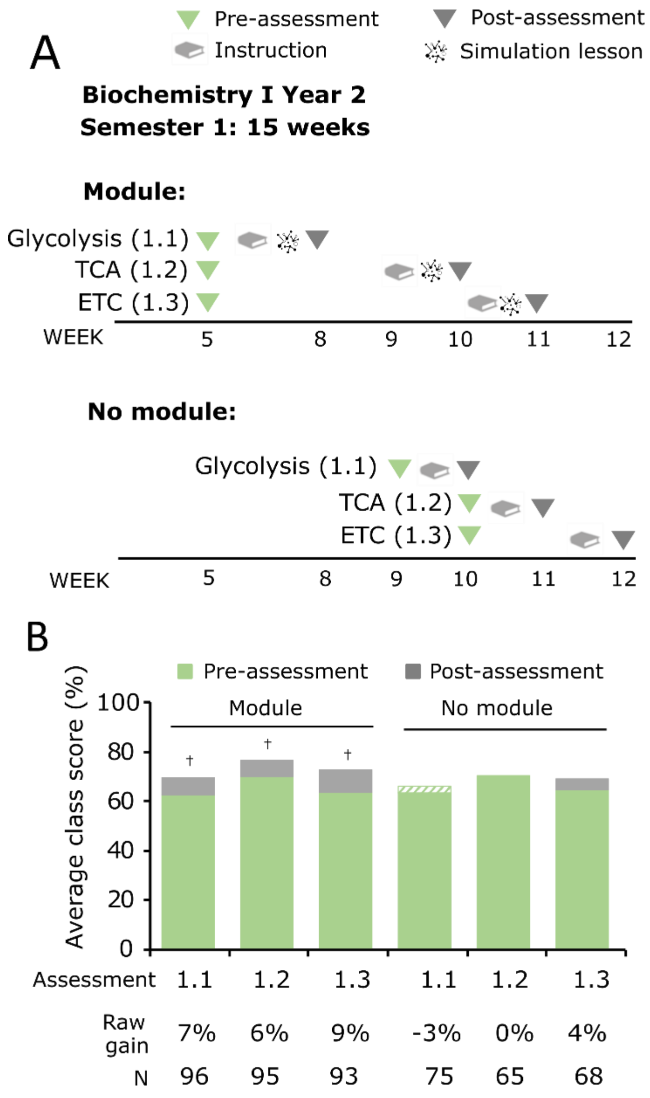


**Fig. S1.** Computational learning modules improve student performance on content assessments for metabolism during Year 2. A) Diagram of the semester for the "Module" (top) and "No module" (bottom) courses of Biochemistry I during Year 2. Assessment and instructional timing for *Regulation of Cellular Respiration* is shown. B) Course average values of the pre-assessment scores (green) and post-assessment scores (grey) were compared between "Module" and "No module" courses for *Cellular Respiration* (Assessment 1.1: *Glycolysis*, Assessment 1.2: *TCA*, Assessment 1.3: *ETC*). Each course was taught by a different instructor (The same instructor from the Year 1 course taught the corresponding Year 2 course). Two-tailed paired t-tests (Supporting Table S2) were used to measure significance for pre- versus post-assessment scores: † indicates $p<0.05$. A green and white striped pattern indicates that the overall post-assessment score was lower than the pre-assessment score.

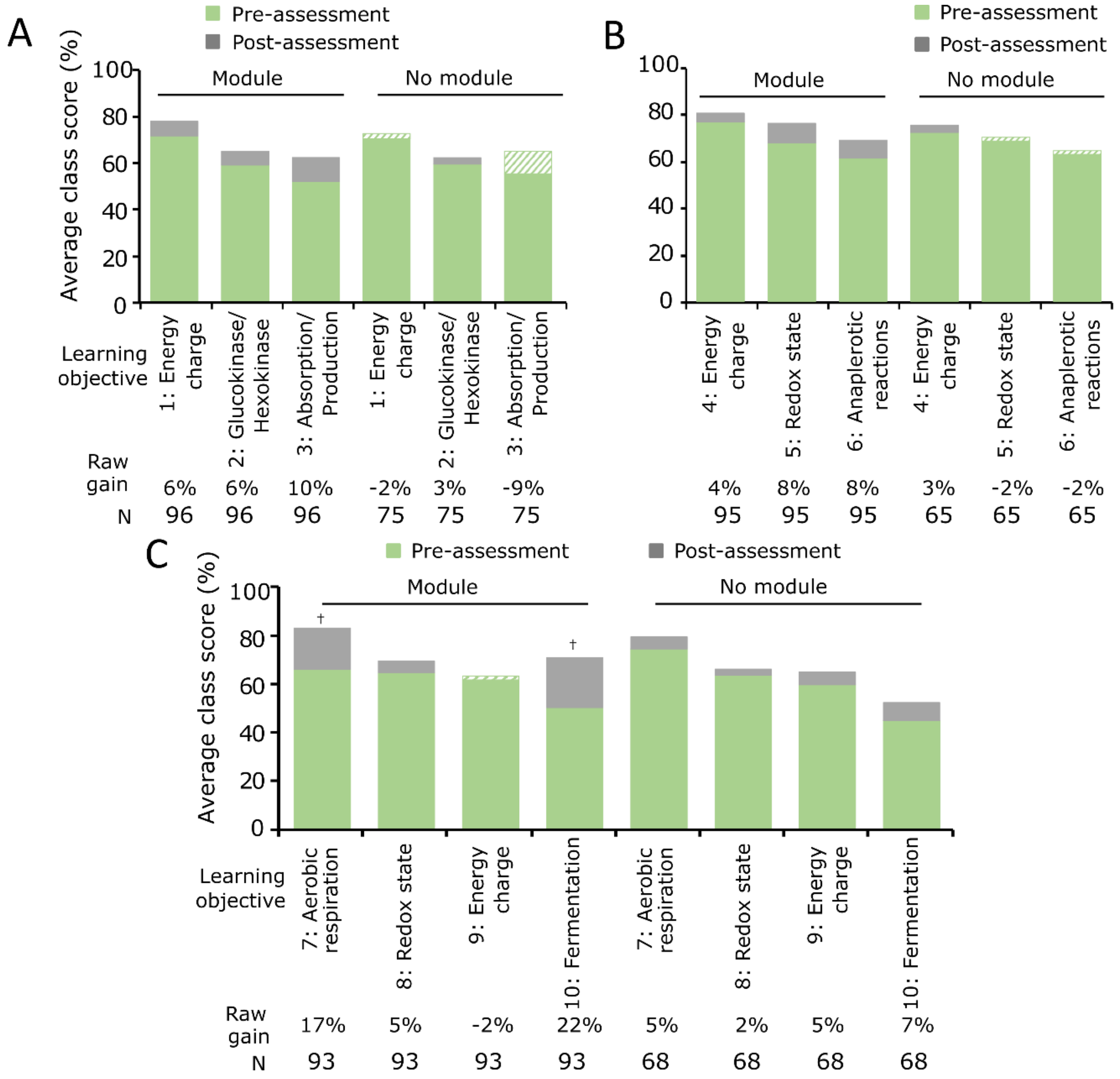


**Fig. S2.** Computational learning modules improve class performance on learning objectives for Cellular Respiration during Year 2. Average class scores of the pre-assessment scores (green) and post-assessment scores (grey) for each stated learning objective in the *Regulation of Cellular Respiration* module during Year 2. A) Assessment 1.1: *Glycolysis*, B) Assessment 1.2: *TCA*, and C) Assessment 1.3: *ETC* were compared for the “Module” and “No module” courses. Each learning objective is numbered, and keywords are provided (refer to Table 1 for detailed objective and corresponding STH level). Two-tailed paired t-tests (Supporting Table S9) were used to measure significance for pre- versus post-assessment scores: † indicates $p<0.05$.

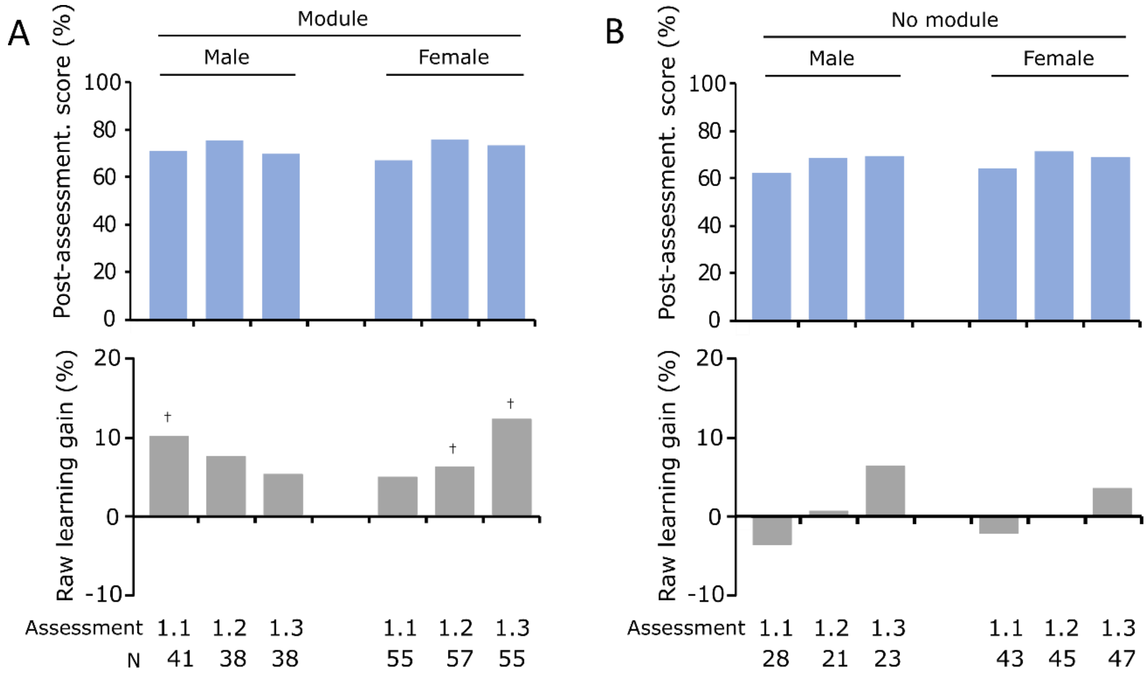


**Fig. S3.** Repeated learning with the computational learning modules may lead to equitable outcomes during Year 2. Course average values of raw learning gains (grey) for students in the A) Biochemistry I "Module" course, and B) Biochemistry I "No module" course. Learning gains were measured for the topic of Regulation of Cellular Respiration using three assessments (Assessment 1.1 = Glycolysis, Assessment 1.2 = TCA, and Assessment 1.3 = ETC). Two-tailed paired t-tests were used to measure significance for pre- versus post-assessment scores: † indicates $p<0.05$.

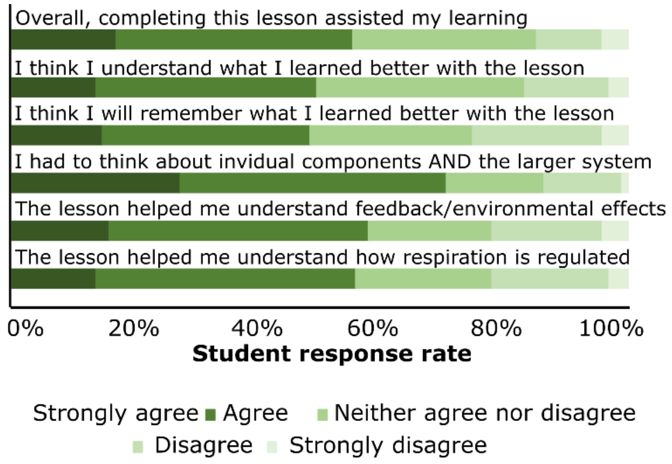


**Fig. S4.** Students valued the modules and report benefits in specific concepts related to systems-thinking. Students in the “Module” courses for Biochemistry I *Regulation of Cellular Respiration* completed a brief survey about their experiences with the module. Results were reported on a five-point Likert scale.

**Table S1.** Class performance on the pre- and post-assessments for each lesson for Biochemistry I.

| | **Module** | | | **No module** | | |
|---|---|---|---|---|---|---|
| **Assessment number** | 1.1: Glycolysis | 1.2: TCA | 1.3: ETC | 1.1: Glycolysis | 1.2: TCA | 1.3: ETC |
| **Pre-assessment class average (%)** | 58.2 | 63.7 | 55.0 | N/A | 61.1 | 62.8 |
| **Pre-assessment SD** | 13.1 | 16.3 | 13.4 | N/A | 16.5 | 15.2 |
| **Post-assessment class average (%)** | 66.7 | 70.3 | 63.7 | N/A | 61.9 | 58.7 |
| **Post-assessment SD** | 17.7 | 18.0 | 18.0 | N/A | 15.3 | 15.6 |
| **Raw learning gain (%)** | 8.5 | 6.6 | 8.8 | N/A | 0.8 | -4.1 |
| **† Pre to post Student's t-test (p-values)** | 0.003 | 0.030 | 0.001 | N/A | 0.773 | 0.095 |

**Table S2.** Class performance on the pre- and post-assessments for each lesson for Biochemistry I during Year 2.

| | **Module** | | | **No module** | | |
|---|---|---|---|---|---|---|
| **Assessment number** | 1.1: Glycolysis | 1.2: TCA | 1.3: ETC | 1.1: Glycolysis | 1.2: TCA | 1.3: ETC |
| **Pre-assessment class average (%)** | 61.5 | 68.8 | 62.4 | 64.9 | 68.9 | 63.4 |
| **Pre-assessment SD** | 17.5 | 16.3 | 18.5 | 13.9 | 13.5 | 16.2 |
| **Post-assessment class average (%)** | 68.3 | 75.3 | 71.4 | 62.2 | 69.1 | 67.8 |
| **Post-assessment SD** | 18.4 | 17.5 | 16.7 | 16.5 | 18.1 | 16.4 |
| **Raw learning gain (%)** | 6.8 | 6.4 | 9.1 | -2.7 | 0.2 | 4.4 |
| **† Pre to post Student's t-test (p-values)** | 0.006 | 0.006 | 0.000 | 0.240 | 0.940 | 0.099 |

**Table S3.** Class performance on the pre- and post-assessments for each lesson for Biochemistry II.

| | **Module: All students** | **Consecutive group** | **Non-consecutive group** |
|---|---|---|---|
| **Assessment number** | 2.1: Purine biosynthesis | 2.1 | 2.1 |
| **Pre-assessment class average (%)** | 57.4 | 57.6 | 57.2 |
| **Pre-assessment SD** | 12.8 | 13.3 | 12.5 |
| **Post-assessment class average (%)** | 61.2 | 64.1 | 58.7 |
| **Post-assessment SD** | 15.8 | 16.7 | 14.7 |
| **Raw learning gain (%)** | 3.8 | 6.5 | 1.5 |
| **† Pre to post Student's t-test (p-values)** | 0.022 | 0.005 | 0.446 |

**Table S4.** Participant demographic profiles for Biochemistry I.

| Demographic variable | Module | | | No module | | | t-statistic | p-value |
|---|---|---|---|---|---|---|---|---|
| | N | Mean | S.D. | N | Mean | S.D. | | |
| **Gender (Male = 0, Female = 1)** | 69 | 0.59 | 0.49 | 73 | 0.68 | 0.47 | 1.122 | 0.264 |
| **Native English Speaker (No = 0, Yes = 1)** | 69 | 0.9 | 0.3 | 73 | 0.97 | 0.16 | 1.79 | 0.076 |
| **Parents' College Education (No = 0, Yes = 1)** | 69 | 0.8 | 0.41 | 73 | 0.78 | 0.42 | -0.236 | 0.814 |
| **Job to Fund College Life (No = 0, Yes = 1)** | 69 | 0.72 | 0.45 | 73 | 0.7 | 0.46 | -0.34 | 0.735 |
| **Cumulative GPA** | 68 | 3.41 | 0.85 | 70 | 3.74 | 0.28 | 3.036 | 0.003 |

**Table S5.** Participant demographic profiles for Biochemistry II.

| Demographic variable | Consecutive group | | | Non-consecutive group | | | t-statistic | p-value |
|---|---|---|---|---|---|---|---|---|
| | N | Mean | S.D. | N | Mean | S.D. | | |
| **Gender (Male = 0, Female = 1)** | 40 | 0.63 | 0.49 | 47 | 0.64 | 0.49 | 0.127 | 0.899 |
| **Native English Speaker (No = 0, Yes = 1)** | 40 | 0.90 | 0.30 | 47 | 0.98 | 0.15 | 1.498 | 0.140 |
| **Parents' College Education (No = 0, Yes = 1)** | 40 | 0.93 | 0.27 | 47 | 0.83 | 0.38 | -1.367 | 0.175 |
| **Job to Fund College Life (No = 0, Yes = 1)** | 40 | 0.75 | 0.44 | 47 | 0.72 | 0.45 | -0.277 | 0.782 |
| **Cumulative GPA** | 40 | 3.64 | 0.39 | 47 | 3.81 | 0.18 | 2.576 | 0.013 |

**Table S6.** One-way ANCOVA results of Module versus No module courses for Biochemistry I.

| Model | Group | N | Unadjusted | | Adjusted | | *F*** | *p-value* | *Partial eta* |
|---|---|---|---|---|---|---|---|---|---|
| | | | M | SD | M* | SE | | | |
| **1.2: TCA** | *Module* | 62 | 0.7 | 0.18 | 0.74 | 0.05 | *(1, 116) 7.443* | *0.007* | *0.060* |
| | *No module* | 62 | 0.62 | 0.15 | 0.65 | 0.05 | | | |
| **1.3: ETC** | *Module* | 55 | 0.64 | 0.18 | 0.63 | 0.05 | *(1, 108) 7.112* | *0.009* | *0.062* |
| | *No module* | 61 | 0.59 | 0.16 | 0.55 | 0.05 | | | |

**Table S7.** One-way ANCOVA results of Consecutive versus Non-consecutive groups for Biochemistry II.

| Model | Group | N | Unadjusted | | Adjusted | | *F*** | *p-value* | *Partial eta* |
|---|---|---|---|---|---|---|---|---|---|
| | | | M | SD | M* | SE | | | |
| **2.1: Purine biosynthesis** | *Consecutive* | 40 | 0.64 | 0.14 | 0.64 | 0.04 | *(1, 79) 8.135* | *0.006* | *0.093* |
| | *Non-consecutive* | 47 | 0.58 | 0.12 | 0.56 | 0.04 | | | |

**Table S8.** Class performance on the pre- and post-assessments for each learning objective for Biochemistry I.

| ***1.1: Glycolysis*** | **Module** | | |
|---|---|---|---|
| **Assessment 1.1 Learning objective** | 1: Energy charge | 2: Glucokinase/ hexokinase | 3: Absorption/ production |
| **Pre-assessment class average (%)** | 60.6 | 50.8 | 59.4 |
| **Pre-assessment SD** | 17.4 | 37.3 | 30.7 |
| **Post-assessment class average (%)** | 68.4 | 70.3 | 58.6 |
| **Post-assessment SD** | 19.5 | 36.4 | 27.5 |
| **Raw learning gain (%)** | 7.8 | 19.5 | -0.8 |
| **† Pre to post Student's t-test (p-values)** | 0.025 | 0.003 | 0.885 |

| ***1.2: TCA cycle*** | **Module** | | | **No module** | | |
|---|---|---|---|---|---|---|
| **Assessment 1.2 Learning objective** | 4: Energy charge | 5: Redox state | 6: Anaplerotic reactions | 4: Energy charge | 5: Redox state | 6: Anaplerotic reactions |
| **Pre-assessment class average (%)** | 72.5 | 65.7 | 52.7 | 74.9 | 66.7 | 49.8 |
| **Pre-assessment SD** | 23.5 | 28.0 | 30.5 | 22.4 | 28.1 | 27.4 |
| **Post-assessment class average (%)** | 76.8 | 77.8 | 57.5 | 70.1 | 70.6 | 50.6 |
| **Post-assessment SD** | 23.1 | 27.8 | 25.5 | 23.3 | 30.6 | 23.3 |
| **Raw learning gain (%)** | 4.3 | 12.1 | 4.8 | -4.8 | 3.9 | 0.9 |
| **† Pre to post Student's t-test (p-values)** | 0.267 | 0.003 | 0.348 | 0.153 | 0.392 | 0.838 |

| ***1.3: ETC and fermentation*** | **Module** | | | | **No module** | | | |
|---|---|---|---|---|---|---|---|---|
| **Assessment 1.2 Learning objective** | 7: Aerobic respiration | 8: Redox state | 9: Energy charge | 10: Ferment. | 7: Aerobic respiration | 8: Redox state | 9: Energy charge | 10: Ferment. |
| **Pre-assessment class average (%)** | 52.2 | 60.8 | 55.0 | 46.7 | 60.3 | 67.1 | 59.0 | 60.3 |
| **Pre-assessment SD** | 24.8 | 19.2 | 31.5 | 50.3 | 24.7 | 23.7 | 29.9 | 49.3 |
| **Post-assessment class average (%)** | 57.2 | 62.1 | 63.3 | 68.3 | 56.0 | 66.2 | 53.8 | 57.7 |
| **Post-assessment SD** | 21.3 | 16.9 | 31.7 | 46.9 | 25.5 | 23.7 | 31.0 | 49.7 |
| **Raw learning gain (%)** | 5.0 | 1.3 | 8.3 | 21.7 | -4.3 | -0.9 | -5.1 | -2.6 |
| **† Pre to post Student's t-test (p-values)** | 0.253 | 0.700 | 0.105 | 0.006 | 0.284 | 0.813 | 0.296 | 0.686 |

**Table S9.** Class performance on the pre- and post-assessments for each learning objective for Biochemistry I during Year 2.

| *1.1: Glycolysis* | Module | | | No module | | |
|---|---|---|---|---|---|---|
| **Assessment 1.1 Learning objective** | 1: Energy charge | 2: Glucokinase/ hexokinase | 3: Absorption/ production | 1: Energy charge | 2: Glucokinase/ hexokinase | 3: Absorption/ production |
| **Pre-assessment class average (%)** | 68.3 | 56.3 | 49.5 | 69.3 | 56.7 | 62.0 |
| **Pre-assessment SD** | 21.5 | 37.2 | 38.7 | 18.4 | 36.1 | 39.3 |
| **Post-assessment class average (%)** | 74.4 | 62.0 | 59.4 | 67.2 | 59.3 | 52.7 |
| **Post-assessment SD** | 21.5 | 38.9 | 37.9 | 17.9 | 36.5 | 37.6 |
| **Raw learning gain (%)** | 6.0 | 5.7 | 9.9 | -2.1 | 2.7 | -9.3 |
| **† Pre to post Student's t-test (p-values)** | 0.054 | 0.299 | 0.053 | 0.465 | 0.645 | 0.094 |

| *1.2: TCA cycle* | Module | | | No module | | |
|---|---|---|---|---|---|---|
| **Assessment 1.2 Learning objective** | 4: Energy charge | 5: Redox state | 6: Anaplerotic reactions | 4: Energy charge | 5: Redox state | 6: Anaplerotic reactions |
| **Pre-assessment class average (%)** | 76.8 | 67.9 | 61.4 | 72.4 | 70.3 | 64.6 |
| **Pre-assessment SD** | 22.8 | 35.7 | 29.3 | 22.7 | 36.4 | 25.1 |
| **Post-assessment class average (%)** | 80.7 | 76.3 | 69.1 | 75.5 | 68.8 | 63.0 |
| **Post-assessment SD** | 21.0 | 35.6 | 26.7 | 24.7 | 33.9 | 27.3 |
| **Raw learning gain (%)** | 3.9 | 8.4 | 7.7 | 3.1 | -1.6 | -1.6 |
| **† Pre to post Student's t-test (p-values)** | 0.212 | 0.077 | 0.051 | 0.458 | 0.805 | 0.728 |

| *1.3: ETC and fermentation* | Module | | | | No module | | | |
|---|---|---|---|---|---|---|---|---|
| **Assessment 1.2 Learning objective** | 7: Aerobic respiration | 8: Redox state | 9: Energy charge | 10: Ferment. | 7: Aerobic respiration | 8: Redox state | 9: Energy charge | 10: Ferment. |
| **Pre-assessment class average (%)** | 65.2 | 63.8 | 62.4 | 49.5 | 73.5 | 62.7 | 58.8 | 44.1 |
| **Pre-assessment SD** | 29.9 | 23.4 | 33.5 | 50.3 | 26.0 | 23.5 | 32.4 | 50.0 |
| **Post-assessment class average (%)** | 82.1 | 68.5 | 60.8 | 69.9 | 78.4 | 65.2 | 64.0 | 51.5 |
| **Post-assessment SD** | 22.8 | 26.2 | 28.4 | 46.1 | 24.9 | 26.0 | 32.1 | 50.3 |
| **Raw learning gain (%)** | 16.8 | 4.7 | -1.6 | 20.4 | 4.9 | 2.5 | 5.1 | 7.4 |
| **† Pre to post Student's t-test (p-values)** | 0.000 | 0.150 | 0.708 | 0.003 | 0.241 | 0.562 | 0.311 | 0.340 |

**Table S10.** Class performance on the pre- and post-assessments for each learning objective for Biochemistry II.

| ***2.1: Purine biosynthesis*** | **Module - All students** | | | |
|---|---|---|---|---|
| **Assessment 2.1 Learning objective** | 1: Components and interactions | 2: Maintain homeostasis | 3: Cellular condition changes | 4: Mutations and disease |
| **Pre-assessment class average (%)** | 55.2 | 62.1 | 61.8 | 55.2 |
| **Pre-assessment SD** | 16.5 | 28.4 | 30.7 | 21.5 |
| **Post-assessment class average (%)** | 62.1 | 59.4 | 60.3 | 61.7 |
| **Post-assessment SD** | 23.7 | 31.5 | 26.3 | 21.6 |
| **Raw learning gain (%)** | 6.9 | -2.7 | -1.4 | 6.5 |
| **† Pre to post Student's t-test (p-values)** | 0.012 | 0.528 | 0.725 | 0.033 |
| | **Consecutive group** | | | |
| **Assessment 2.1 Learning objective** | 1: Components and interactions | 2: Maintain homeostasis | 3: Cellular condition changes | 4: Mutations and disease |
| **Pre-assessment class average (%)** | 58.2 | 60.8 | 61.3 | 52.9 |
| **Pre-assessment SD** | 16.3 | 29.1 | 29.4 | 22.6 |
| **Post-assessment class average (%)** | 65.0 | 60.8 | 64.4 | 64.6 |
| **Post-assessment SD** | 24.8 | 30.1 | 23.9 | 19.3 |
| **Raw learning gain (%)** | 6.8 | 0.0 | 3.1 | 11.7 |
| **† Pre to post Student's t-test (p-values)** | 0.071 | 1.000 | 0.515 | 0.003 |
| | **Non-consecutive group** | | | |
| **Assessment 2.1 Learning objective** | 1: Components and interactions | 2: Maintain homeostasis | 3: Cellular condition changes | 4: Mutations and disease |
| **Pre-assessment class average (%)** | 52.6 | 63.1 | 62.2 | 57.1 |
| **Pre-assessment SD** | 16.3 | 28.0 | 32.1 | 20.5 |
| **Post-assessment class average (%)** | 59.6 | 58.2 | 56.9 | 59.2 |
| **Post-assessment SD** | 22.7 | 32.9 | 27.9 | 23.3 |
| **Raw learning gain (%)** | 7.0 | -5.0 | -5.3 | 2.1 |
| **† Pre to post Student's t-test (p-values)** | 0.078 | 0.425 | 0.407 | 0.642 |

**Table S11.** One-way ANCOVA results of gender groups for Biochemistry I.

| Model | Group | N | Unadjusted | | Adjusted | | *F*** | *p-value* | *Partial eta* |
|---|---|---|---|---|---|---|---|---|---|
| | | | M | SD | M* | SE | | | |
| **1.2: TCA** | *Module and male* | 25 | 0.74 | 0.19 | 0.75 | 0.05 | *(3, 115) 3.021* | *0.033* | *0.073* |
| | *Module and female* | 37 | 0.68 | 0.18 | 0.7 | 0.04 | | | |
| | *No module and male* | 21 | 0.63 | 0.12 | 0.63 | 0.05 | | | |
| | *No module and female* | 41 | 0.62 | 0.17 | 0.63 | 0.04 | | | |
| **1.3: ETC** | *Module and male* | 22 | 0.64 | 0.17 | 0.6 | 0.05 | *(3, 107) 2.462* | *0.067* | *0.065* |
| | *Module and female* | 33 | 0.64 | 0.19 | 0.64 | 0.05 | | | |
| | *No module and male* | 20 | 0.6 | 0.21 | 0.54 | 0.05 | | | |
| | *No module and female* | 41 | 0.58 | 0.12 | 0.54 | 0.04 | | | |

**File S1**

Using our computational learning modules, instructors can employ different adoption approaches to meet their specific course needs and teaching strategies. Our experience indicates that students who have never used the models before and are first exposed to them when learning about unfamiliar biochemistry content may be overwhelmed in the beginning. Sometimes the situation of concurrently introducing a new teaching approach and unfamiliar content is unavoidable. We believe that instructors can mediate student difficulties using a variety of strategies, including using a guided-instruction (described second) or laboratory-linked approach (described third).

First, instructors can integrate the modules as we did, using the course slides below as a guideline. If all three parts of the *Regulation of Cellular Respiration* module will be used, we recommend that students complete them in the order presented in the manuscript. However, instructors may decide to focus only on one part if they provide the appropriate support for students. Similarly, the *Regulation of Purine Biosynthesis* module covers different aspects of this system's regulation, and it may be possible to use only a subset of the module activities. Instructors could also incorporate our assessment questions into their regularly scheduled exams or quizzes to reduce assessment fatigue.

Second, instructors can follow the guided-instruction approach by fully discussing the system and how the components fit together before introducing the models in class. Instructors could then introduce the models and module questions during lectures where students discuss and respond to the questions and report back to the instructor during class (either as whole-class group feedback or clicker responses). With this alternative approach, the instructor will be manipulating the model and dealing with possible technological issues while the students are engaged conceptually with the material using group discussion as opposed to being focused on discussions about modeling instructions or troubleshooting. Once the instructors and students are comfortable with the new approach, the instructor may then ask students to complete subsequent parts of the module on their own in class or as homework.

Third, instructors could integrate the modules as part of a hands-on laboratory experiences. Using this approach, instructors can ask student to make predictions using the models that can then be tested during the laboratory. Students could also use the models to design their laboratory experiments before testing their predictions.

Finally, our modules may be ideal for instructors who are using online or blended courses where students complete the module completely as homework. The approach we used in our courses required that significant portions of the modules were completed as homework, so we believe that students will be successful with this approach

#### Timing

These slides should be introduced after students have completed the glycolysis portion of the *Regulation of Cellular Respiration* module

#### Prerequisite Knowledge and suggestions for incorporation

Introduce after discussing the following topics:

1) Steps in glycolysis
2) Regulation of glycolysis

#### Glycolysis: Phases

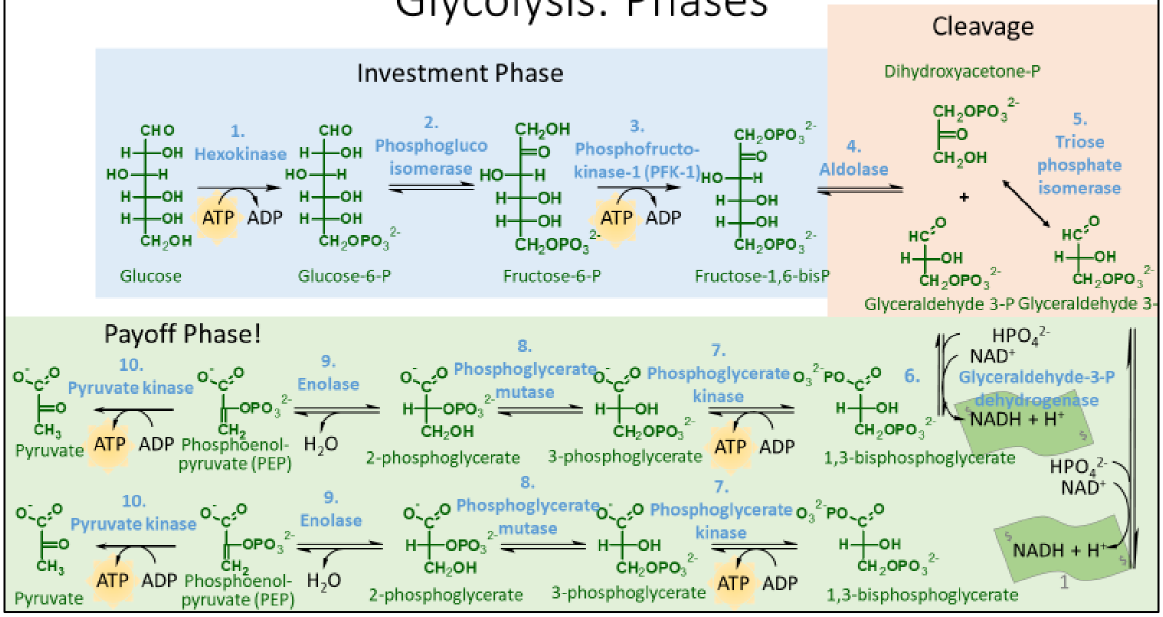

### Regulation

Consider the changes from $\Delta G^{\circ\prime}$ to $\Delta G$ in erythrocytes:

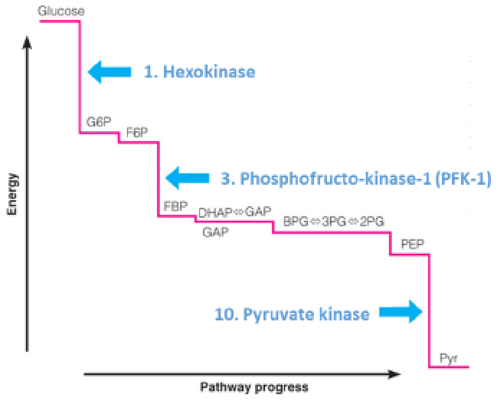


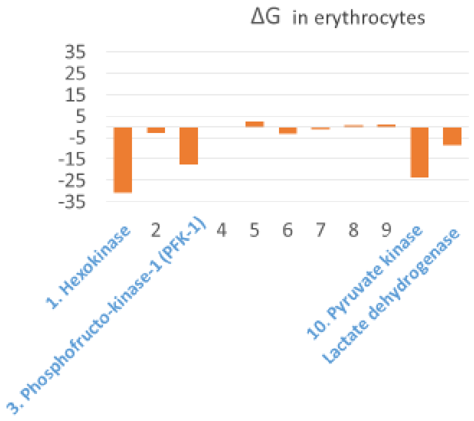


### Glycolysis regulation summary

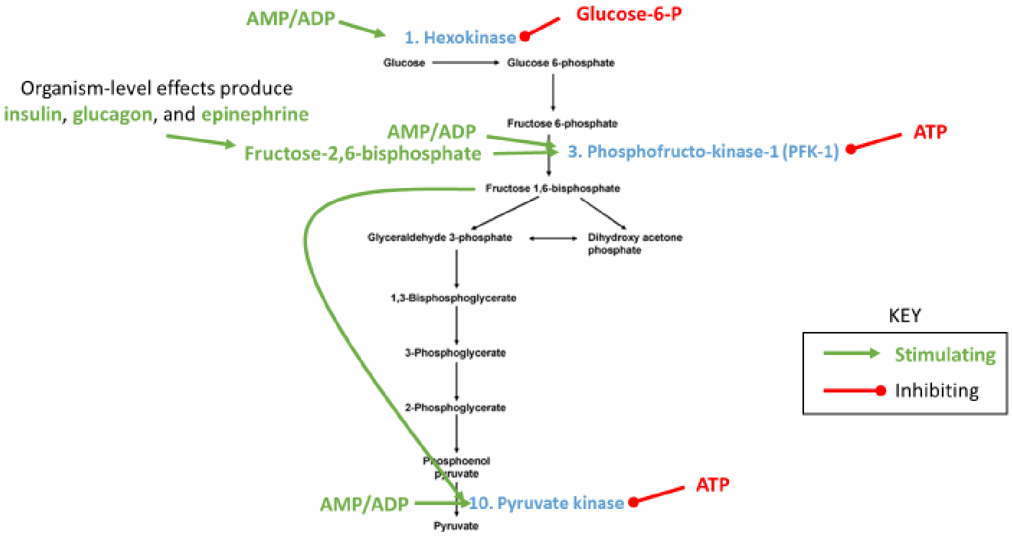

### Hexokinase regulation

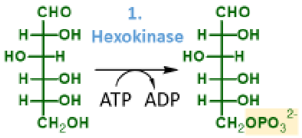


Hexokinase can be considered the "**committed step**" of **glucose entering the cell**.

Negatively regulated by **Glucose-6-P**

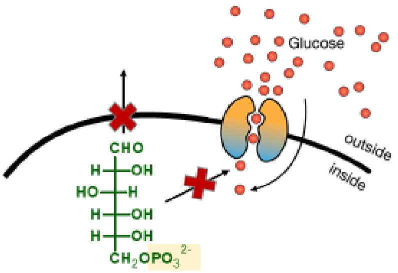


This is NOT the "committed step" of glycolysis.
i.e., glucose-6-P is used in other pathways.

### Hexokinase vs Glucokinase regulation: Modeling

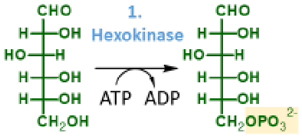


Not all tissues require equal amounts of glucose.

Only two tissues store significant amounts of glycogen:

Liver & muscle

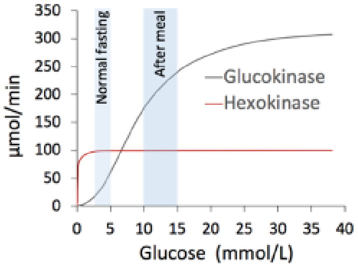


The **liver stores glucose as glycogen** primarily to **maintain blood sugar** levels between meals. This is very important for the brain.

Hexokinase is feedback inhibited by **Glucose-6-P**
Glucokinase is not.

Why? We will use a model to answer this.

### Phosphofructokinase regulation

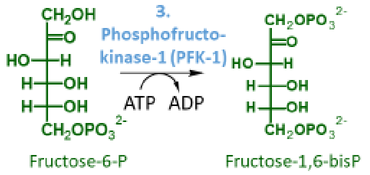


Phosphofructokinase is the **committed step** of glycolysis

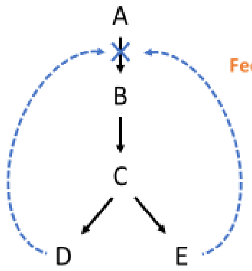


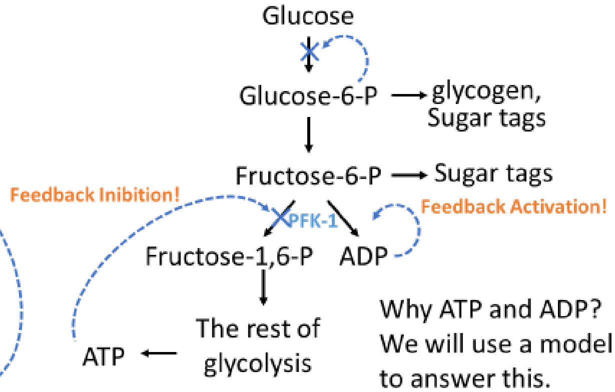


Why ATP and ADP? We will use a model to answer this.

### Pyruvate kinase regulation

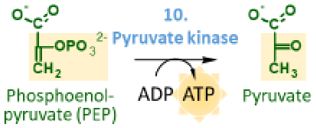


Without the "pull" of pyruvate kinase, there is no way to make reactions 4 -9 stable. It will also effectively stop glycolysis.

Why regulate the last step of the pathway?

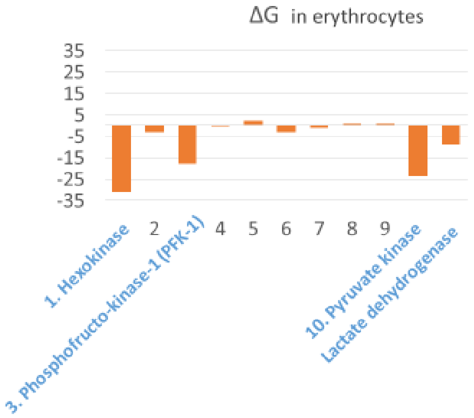

#### Clicker Question

What parts of metabolism do *not* need to be included to build an accurate metabolic model?

A. Reversible reactions
B. Reactions with *no* allosteric control
C. Reactions not relevant to the question asked of the model
D. All of the above

#### Questions we need a glycolysis model to answer:

1. Why are Hexokinase and Glucokinase regulated differently?
2. Why is phosphofructokinase regulated by ATP *and* ADP?
3. Why is pyruvate kinase *also* regulated?
4. Why is pyruvate kinase *also* regulated by ATP *and* ADP?

### In-class activity: Model glycolysis regulation

https://learn.cellcollective.org/#

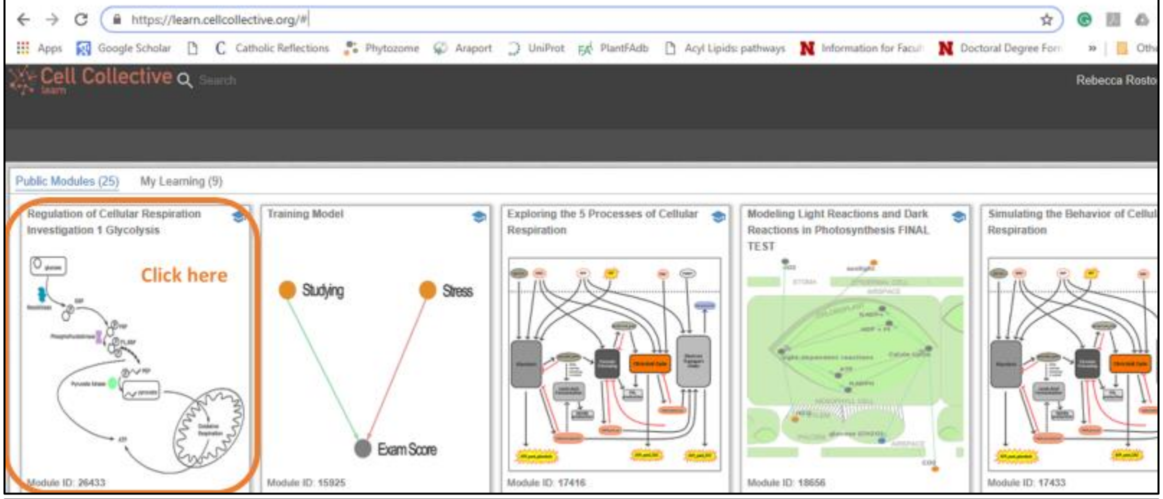


### In-class activity: Model glycolysis regulation

https://learn.cellcollective.org/#

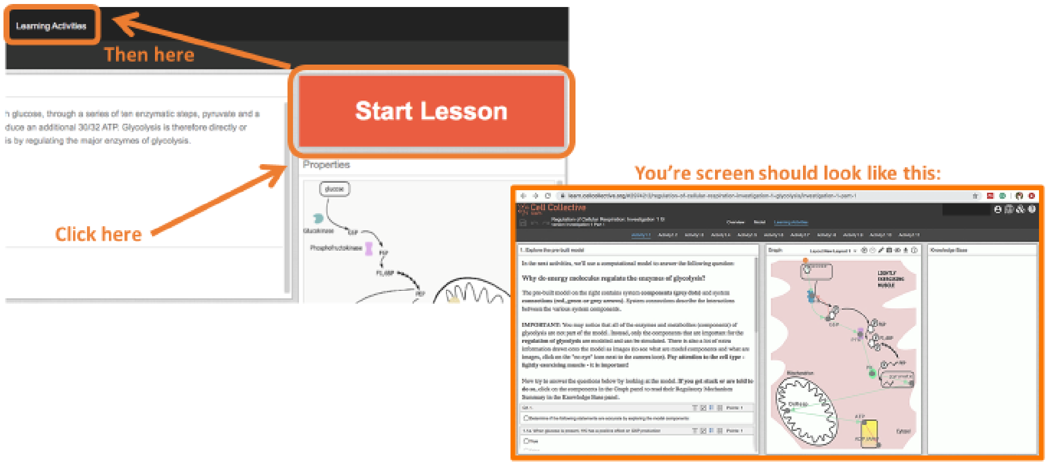

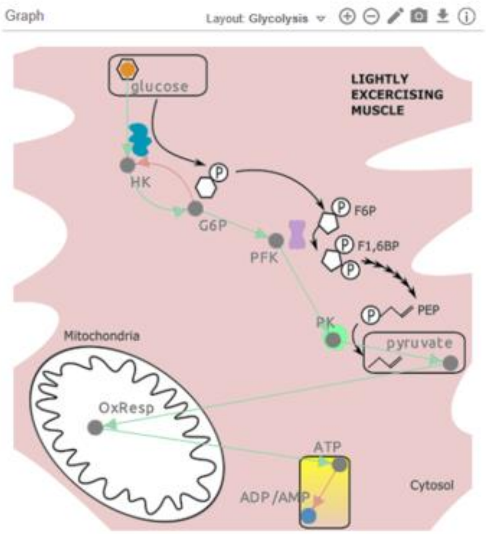


Some metabolites (F1,6BP and PEP) are shown as images, but are not system components and cannot be simulated.

**You can only change dots (nodes) and arrows (edges).**

**Pay attention to the cell type - lightly exercising muscle – these results are not true for all cell types.**

Clicker Question: Click in with "A" when you have reached the thought question on Activity 1.2

Thought Question: A lightly exercising muscle will have a relatively constant need for ATP synthesis…

#### Activity 1.2 Class Discussion Question

Do you think that the rate of glycolysis should change as the [glucose] changes? Why?

"Yes"es reasons:

"No"s reasons:

#### How to…

##### Change the simulation speed

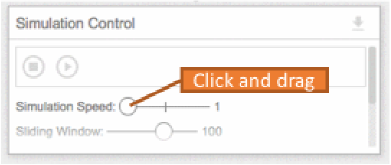


##### Undo/Save

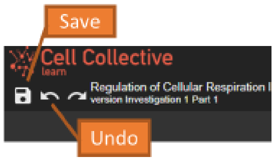


##### Add connectors

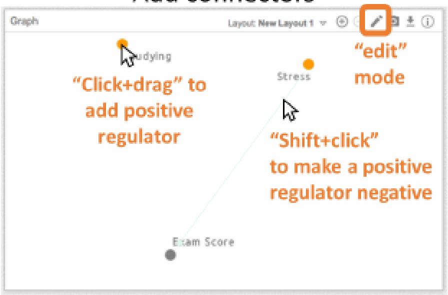


##### Select components to view

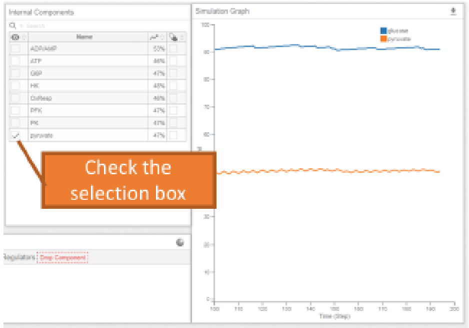


##### Remove connectors

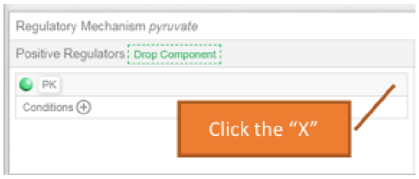

#### Timing

These slides should be introduced after students have completed the glycolysis portion of the *Regulation of Cellular Respiration* module

#### Prerequisite Knowledge and suggestions for incorporation

Introduce after discussing the following topics:

1) Steps in glycolysis
2) Regulation of glycolysis

### Last week: Glycolysis

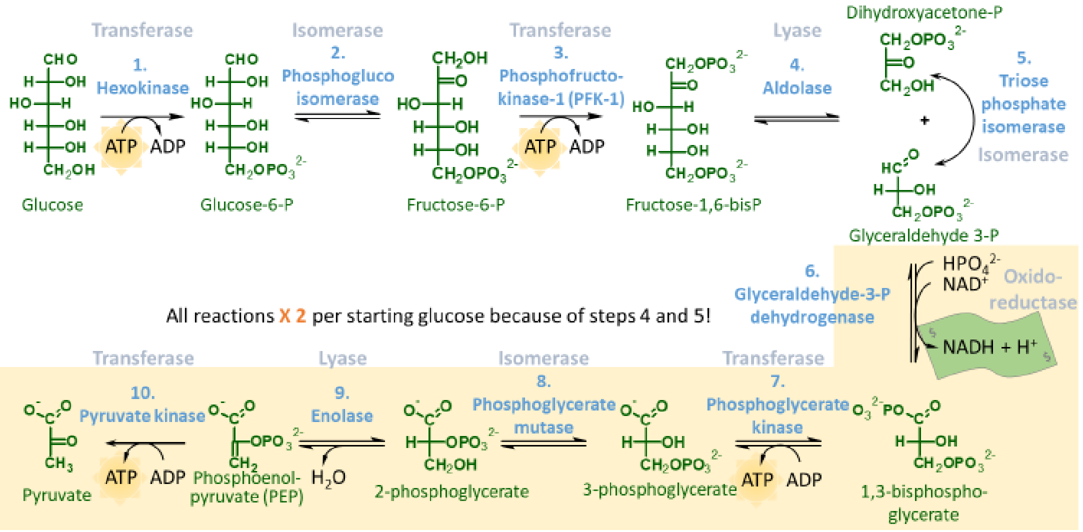

#### Questions we need a glycolysis model to answer:

1. Why are Hexokinase and Glucokinase regulated differently?
2. Why is phosphofructokinase regulated by ATP *and* ADP?
3. Why is pyruvate kinase *also* regulated?
4. Why is pyruvate kinase *also* regulated by ATP *and* ADP?

#### Regulation of Glycolysis

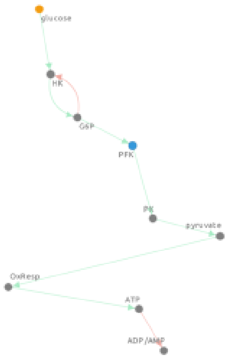


High glucose and no regulation of Phosphofructokinase or Pyruvate kinase

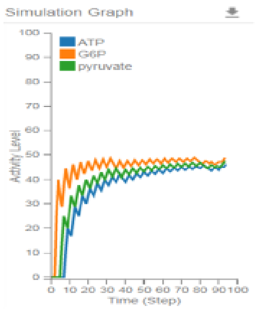


Low glucose and no regulation of Phosphofructokinase or Pyruvate kinase

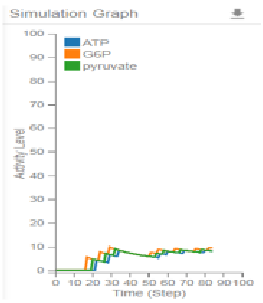


##### Clicker Question

Notice the ATP levels. Knowing that ATP:ADP ratio was important for making ΔG favorable during glycolysis, *predict if this cell will survive* when glucose is low.

A. Yes

B. No or not well

### Regulation of Glycolysis

#### Clicker Question

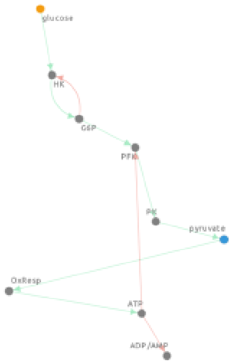


**High glucose** and **ATP negative regulation** of **Phosphofructokinase** only

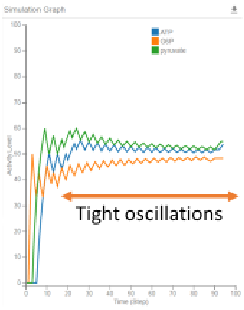


**Low glucose** and **ATP negative regulation** of **Phosphofructokinase** only

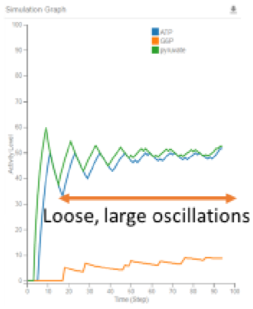


Is simple regulation of one enzyme enough to help maintain cellular ATP levels?

A. Yes

B. No or not well

### Regulation of Glycolysis

#### Clicker Question

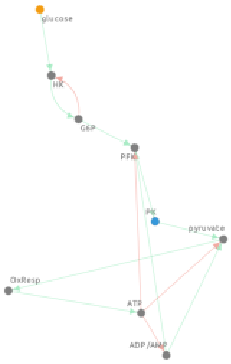


**High glucose** and **ATP/ADP regulation** of **Phosphofructokinase** and **Pyruvate kinase**

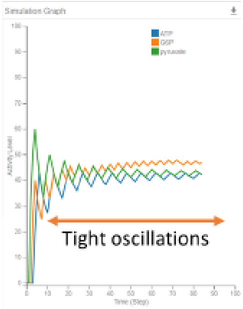


**Low glucose** and **ATP/ADP regulation** of **Phosphofructokinase** and **Pyruvate kinase**

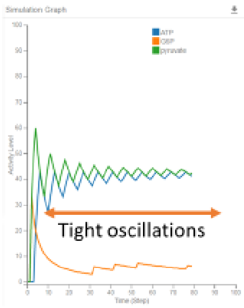


Does regulation of multiple enzymes improve cellular ATP regulation over that of one enzyme?

A. Yes

B. No or not well

### Regulation of Glycolysis

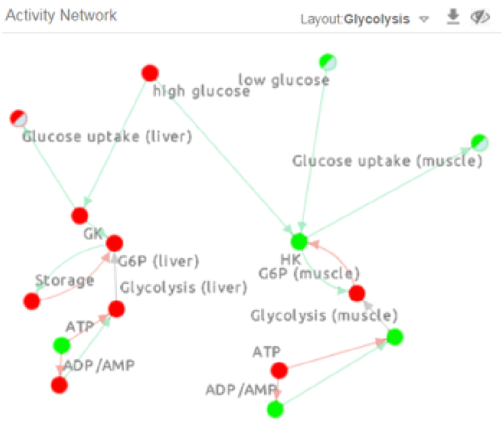


#### Clicker Question

When glucose is low, which tissues take it up?

A. Muscle

B. Liver

C. A and B

### Regulation of Glycolysis

**High glucose** **Low glucose**

Monitoring Glycolysis Output (ATP production)

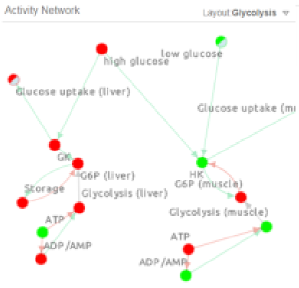


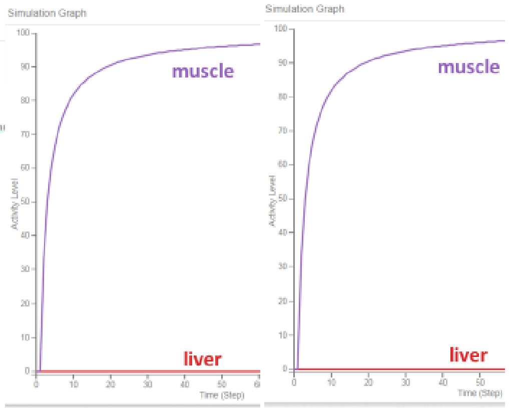


#### Clicker Question

Does the glycolysis level in either tissue depend on the uptake of glucose?

A. Yes

B. No or not much

#### Timing

These slides should be introduced before students have completed the TCA portion of the *Regulation of Cellular Respiration* module

They could be introduced during the same class as the previous slide set

#### Prerequisite Knowledge and suggestions for incorporation

Introduce concurrently with the following topics:

1) Steps in the TCA cycle
2) Regulation of the TCA cycle

#### Regulation of the TCA cycle

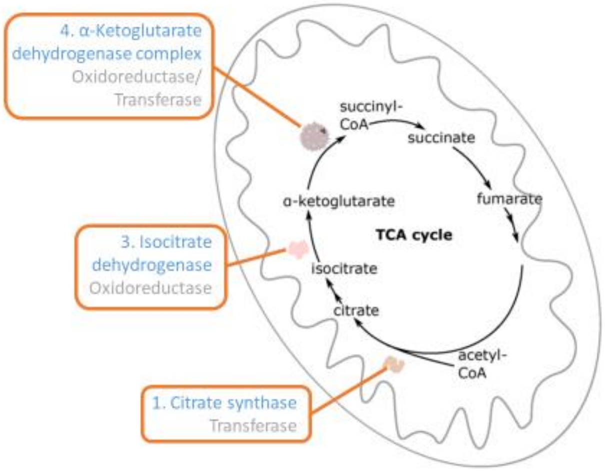

### Regulation of the TCA cycle

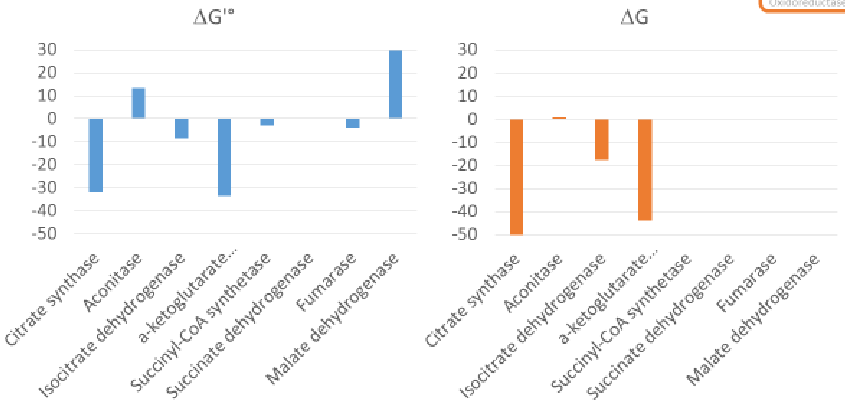


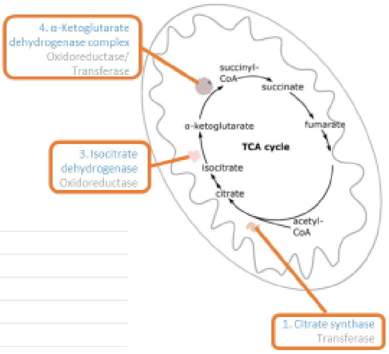


### Regulation of the PDC & TCA cycle

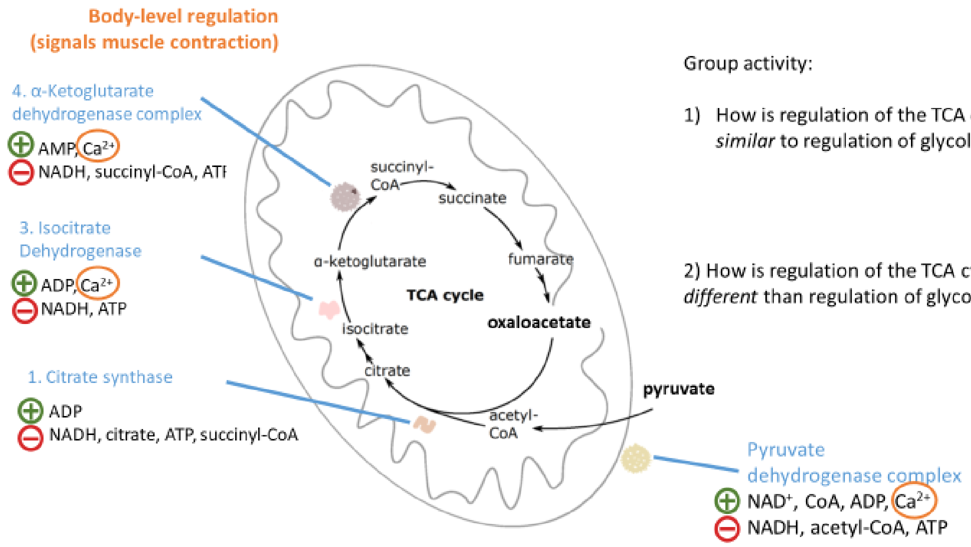


Group activity:

1) How is regulation of the TCA cycle *similar* to regulation of glycolysis?

2) How is regulation of the TCA cycle *different* than regulation of glycolysis?

### Regulation of the PDC & TCA cycle

**Body-level regulation (signals muscle contraction)**

Two themes:
**product inhibition** and **substrate availability**

Model:

How will anaplerotic reactions help maintain TCA function?

Scenario: Cells need energy to build *and* amino acids to build with.

4. α-Ketoglutarate dehydrogenase complex
⊕ AMP, $Ca^{2+}$
⊖ NADH, succinyl-CoA, ATP

3. Isocitrate Dehydrogenase
⊕ ADP, $Ca^{2+}$
⊖ NADH, ATP

1. Citrate synthase
⊕ ADP
⊖ NADH, citrate, ATP, succinyl-CoA

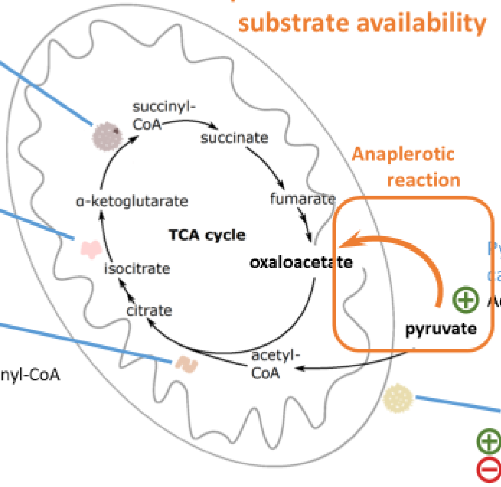


Pyruvate carboxylase
⊕ Acetyl-CoA

Pyruvate dehydrogenase complex
⊕ $NAD^+$, CoA, ADP, $Ca^{2+}$
⊖ NADH, acetyl-CoA, ATP

### Cell Collective Homework Assignment!

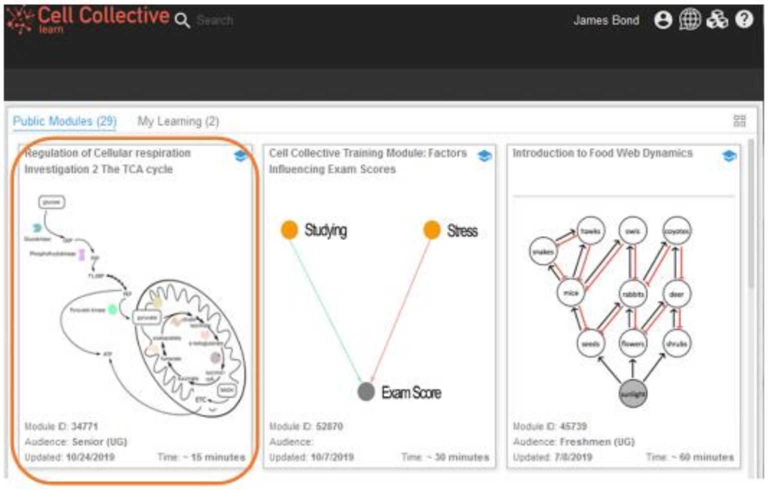

### The TCA cycle is not only used for energy!

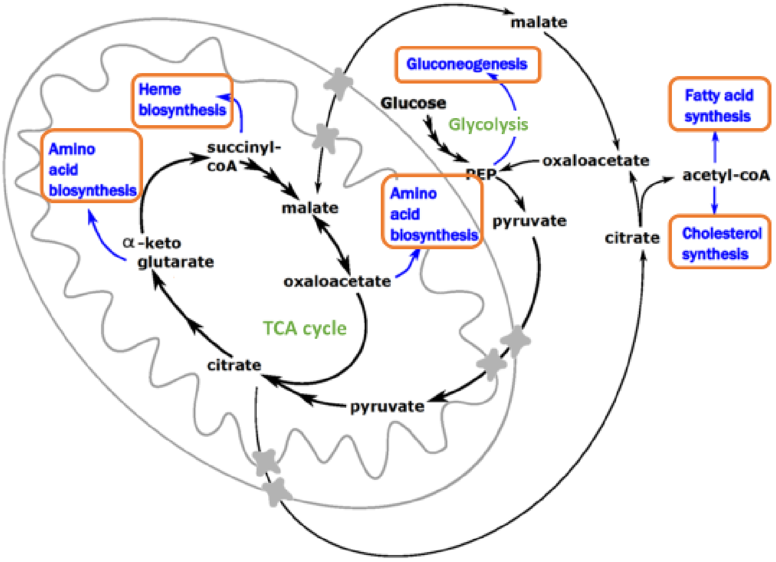


### The TCA cycle is not only used for energy!

These are called anaplerotic reactions

They are filling up intermediates of a metabolic pathway

Ancient Greek: anaplerosis
ἀνά= 'up' and πληρόω= 'to fill'.

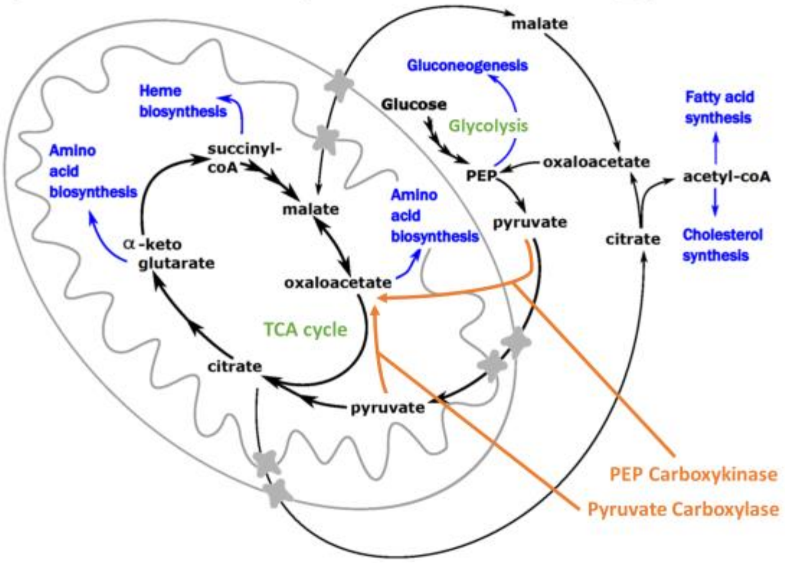

#### Timing

These slides should be introduced before students have completed the ETC portion of the *Regulation of Cellular Respiration* module

#### Prerequisite Knowledge and suggestions for incorporation

Introduce after discussing the following topics:

Fermentation

Introduce concurrently with the following topics:

1) Steps in the ETC
2) Regulation of the ETC

### Homework assignment:

Testing-type questions: predict what happens to respiration when…

Simulation Results Table 3.1: Are the following processes active (Y/N)?

| Environmental/cellular condition → | O2 present, No LDH | No O2 or LDH | LDH present, No O2 |
|---|---|---|---|
| Pyruvate (glycolysis) | Enter Text | Enter Text | Enter Text |
| Lactate (fermentation) | Enter Text | Enter Text | Enter Text |
| CO2 (TCA cycle) | Enter Text | Enter Text | Enter Text |
| O2 consumption (ETC) | Enter Text | Enter Text | Enter Text |

Simulation Results Table 3.2: Is ATP produced by the processes (Y/N)?

| Environmental/cellular condition → | O2 present, No LDH | No O2 or LDH | LDH present, No O2 |
|---|---|---|---|
| 2 ATP (glycolysis) | Enter Text | Enter Text | Enter Text |
| 30/32 ATP (TCA and ETC) | Enter Text | Enter Text | Enter Text |

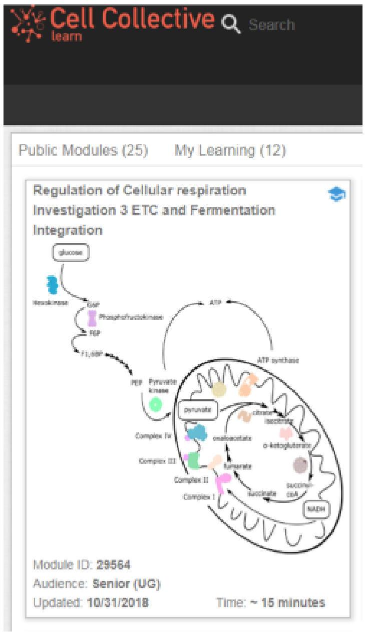

#### Timing

These slides should be introduced after students have completed the ETC portion of the *Regulation of Cellular Respiration* module

#### Prerequisite Knowledge and suggestions for incorporation

Introduce after discussing the following topics:

1) Fermentation
2) Steps in the ETC
3) Regulation of the ETC

### Homework on Respiration using modeling

With $O_2$, energy is effectively supplied | Without fermentation, $O_2$ is needed. | Without $O_2$, more glycolysis is needed.

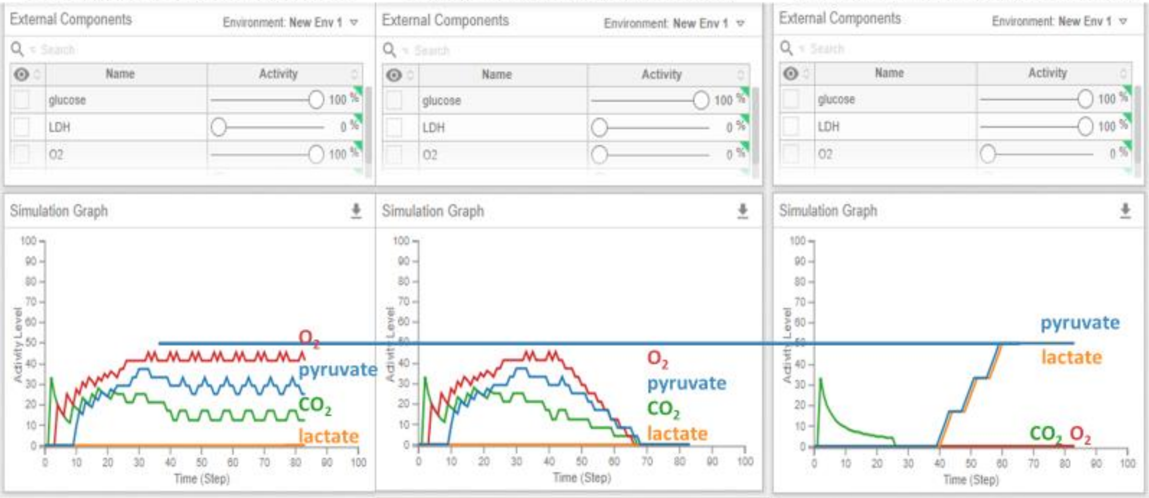

Clicker question:

### Homework on Respiration using modeling

Why does flux (numbers of substrates/intermediates) through glycolysis increase in the absence of $O_2$?

A. Need for $NAD^+$ regeneration
B. ETC is working harder
C. Need for ATP
D. A and B
E. All of the above

### ATP yield of 1 glucose:

| Pathway | | ATP | Reductant | ATP equivalents |
|---|---|---|---|---|
| Glycolysis | 1 Glucose → 2 Pyruvate | **2 ATP** | **2 NADH*** | **2 + 5*** |
| Pyruvate Dehydrogenase | 2 Pyruvate → 2 Acetyl-CoA + 2 $CO_2$ | | **2 NADH** | **5** |
| TCA Cycle | 2 Acetyl-CoA → 4 $CO_2$ | **2 GTP** | **6 NADH**<br>**2 $FADH_2$** | **2 + 15**<br>**3** |
| | 1 NADH = 2.5 ATP<br>1 $FADH_2$ = 1.5 ATP<br>1 GTP = 1 ATP | | | **32 ATP** |

*30 ATP if glycerol-3-P shuttle

If we labeled all carbons on the glucose, adding only 1 labeled glucose, into a cell filled with unlabeled glucose, where are our labels now?

### Cell choices

- PDC/TCA/ETC/shuttle

  OR

- Fermentation

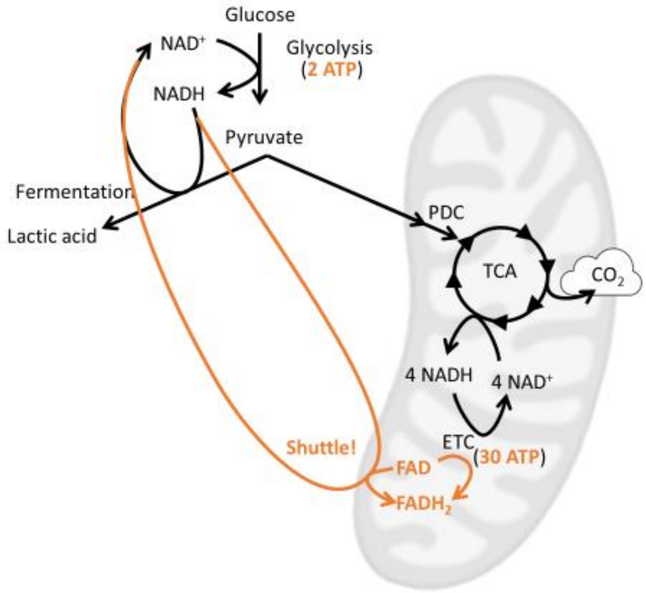


### NOT Cell Choices (creates dead cells…)

- Let the ADP/ATP ratio drop
- Let the NAD$^+$/NADH ratio drop

These are required for homeostasis

The cytosolic NAD$^+$/NADH ratio is maintained.

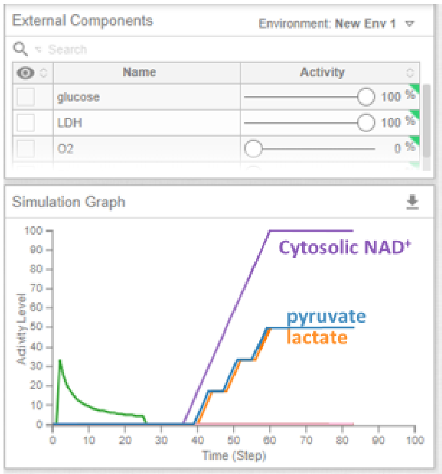

### Modeling Respiration & Exercise

Without exercise, TCA is working constantly

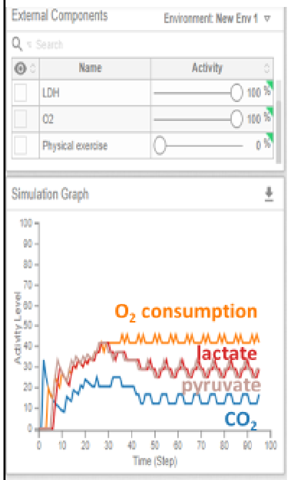


With exercise, TCA is begins to increase

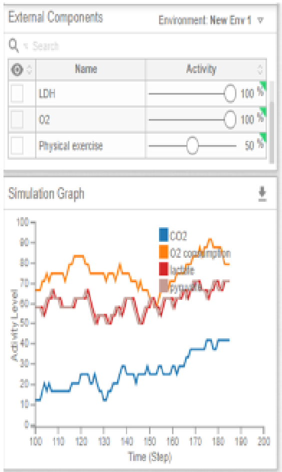


As exercise begins to make $O_2$ limiting, TCA use drops

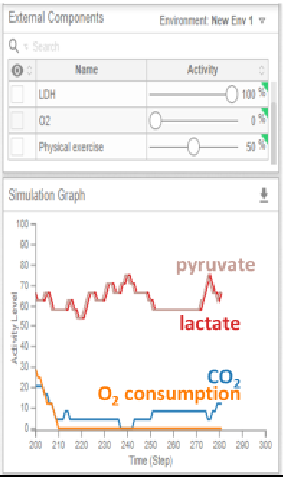


What would you expect about cytosolic [NAD⁺] and [ATP] before and during hard exercise?

A. Same

B. Different

Use caution when interpreting ATP from the model!

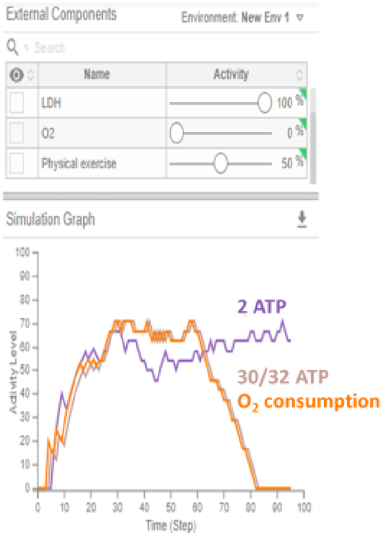

**File S3**

### Regulation of Cellular Respiration and Fermentation Instructor Guides

#### Part 1: Glycolysis

(Module ID: 29742 at https://cellcollective.org)

The diagram below shows the components of cellular respiration that are covered in this module:

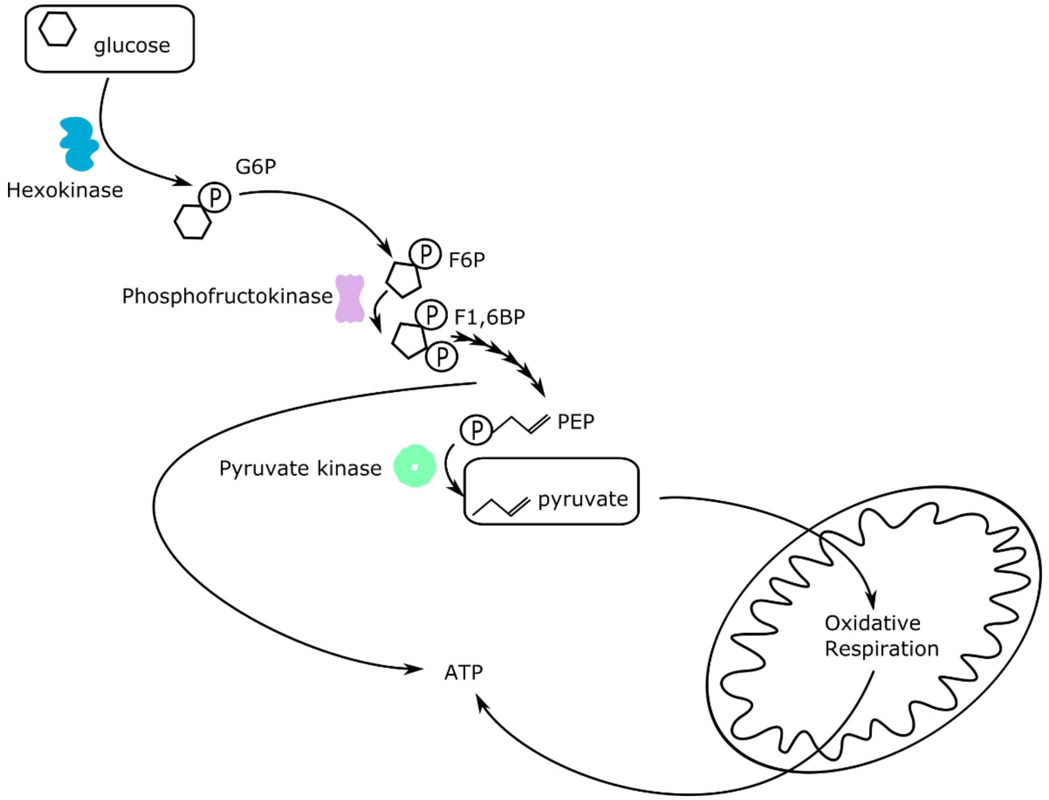

The goal of the first half of the investigation is to introduce students to the importance of **energy charge-based regulation of glycolytic enzymes** to maintain **energy homeostasis**.

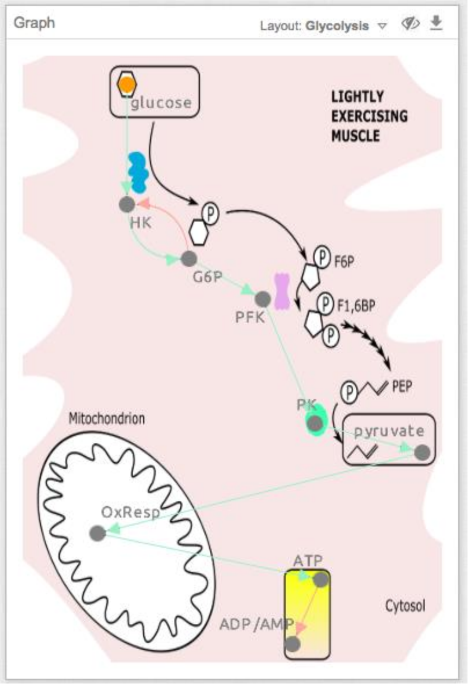


Students are presented with a computational model showing select enzymes and metabolites of glycolysis without most of the known allosteric feedback regulatory connections present (snapshot to the right).

Students are asked to simulate the behavior of the model as is (snapshot to the right) and then to add negative allosteric feedback relationships from ATP to each of the three enzymes that catalyze an irreversible step of glycolysis. They simulate the behavior of the model as they add each of these regulatory connections, tabulate the simulation results. Finally, they evaluate their simulation results before and after adding the negative allosteric feedback relationships. They then follow the same procedures for the positive allosteric feedback regulatory relationships from ADP/AMP to each of the three glycolytic enzymes.

Throughout the investigation, students are asked to reason about how these regulatory connections will affect the entire organism.

#### How instructors can help

Before students start the module:

1. Remind them to add the module to My Learning (clicked ADD TO MY LEARNING), via the Overview page of the module. This will enable the module to be edited.
2. Direct them to the Learning Activities page in Cell Collective.
3. Ask students to confirm that the model is in “edit” mode (this should be the default and is indicated by a “pencil” icon, however, if the model is in “view” mode, students can click the “eye” icon within the Graph panel to change it to “edit” mode).
4. Remind students to:

a. To draw a connection (arrow), click the starting component, drag, and release the mouse over the component you want the arrowhead to land on.

b. To delete an arrow, highlight it, then press delete (fn+delete for mac).

c. To toggle an arrow from positive to negative: (1) deselect everything! (2) press and hold shift while you next click on the arrow that you want to change. Click somewhere else to see the effect.

Although this is pointed out explicitly in the lesson, it may still be important to remind students to pay attention to the cell type and oxygenation status that is represented by each model. Students will be asked many conceptual questions throughout the activities that will require them to critically assess the purpose of homeostasis in the organism. It may be helpful to remind them to draw from previous biology experience or discuss their thinking with a partner/instructor.

#### Model connection building/simulation review

1. ATP negatively regulates glycolysis and reduces the flux through the glycolysis pathway.
2. ADP positively regulates glycolysis and increases the flux through glycolysis.
3. Energy charge-based regulation by ATP and ADP/AMP ensures that the cell always has sufficient energy supply (maintains homeostasis!) regardless of how much glucose is available.

The goal of the second half of the investigation is to have students evaluate the **differences in glucokinase (GK) and hexokinase (HK) kinetics** as a partial explanation of **tissue-specific differences in glucose absorption**.

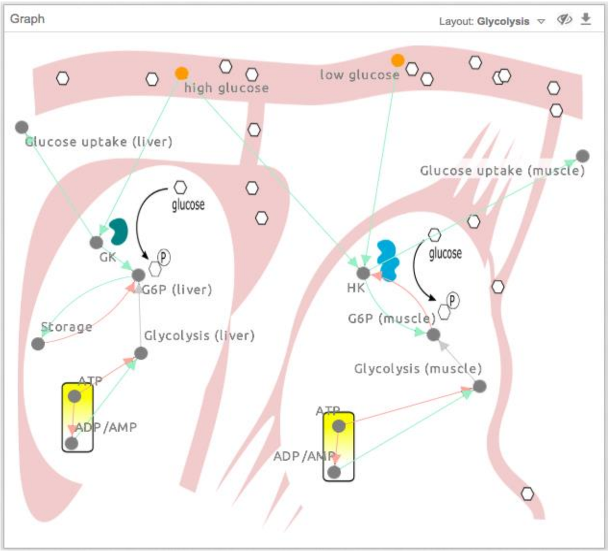


Using a kinetic diagram (not shown), students are asked to predict how glucokinase (GK) and hexokinase (HK) differentially affect glucose absorption in different tissues. Students are then presented with a computational model showing select components and feedback regulatory connections already present in two cell types (snapshot to the right). Through simulation, students discover that GK and HK activity do not determine pyruvate production; instead, the activities of PFK and PK (first part of the investigation) are the major determinants of pyruvate production.

#### How instructors can help

Although this is covered explicitly in the lesson through direct questioning, students may still struggle to connect the kinetic diagram with the simulation output. Students may also still require additional support as they reason through the system.

#### Model connection building/simulation review (continued from the first half of the investigation)

4. Glucokinase and hexokinase determine whether glucose uptake will occur in liver or muscle cells in response to glucose availability.
5. Glucokinase can be active and take up a lot of glucose even if glycolysis stays low because glucose can be stored.
6. Regulation of glucokinase (GK) and hexokinase (HK) is not the major determinant of pyruvate production (glycolysis). Instead, regulation of phosphofructokinase (PFK) and pyruvate kinase (PK) is the major determinant of pyruvate production (glycolysis).

#### Part 2: TCA

(Module ID: 34771 at https://cellcollective.org)

The diagram below shows the components of cellular respirations that are covered in this module:

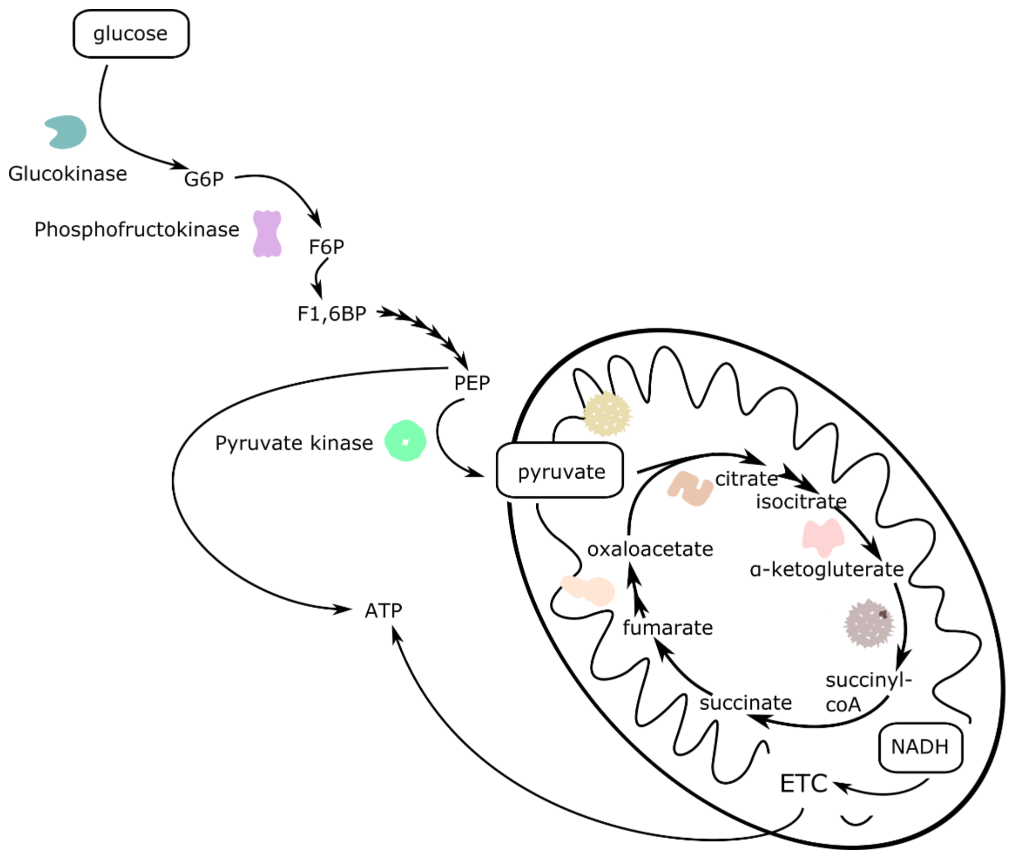

The goal of the first half of the investigation is to introduce students to the importance of **allosteric feedback regulation of TCA enzymes by NADH and energy molecules** to maintain **redox balance (favorable cellular conditions)** and **how these connections affect metabolites produced from glycolysis**.

Students are presented with a computational model showing select enzymes and metabolites of glycolysis and the TCA cycle. In this model, the known feedback regulatory connections to TCA cycle enzymes are not yet present (snapshot to the right - top). Students are asked to simulate the behavior of the model as is and record the results.

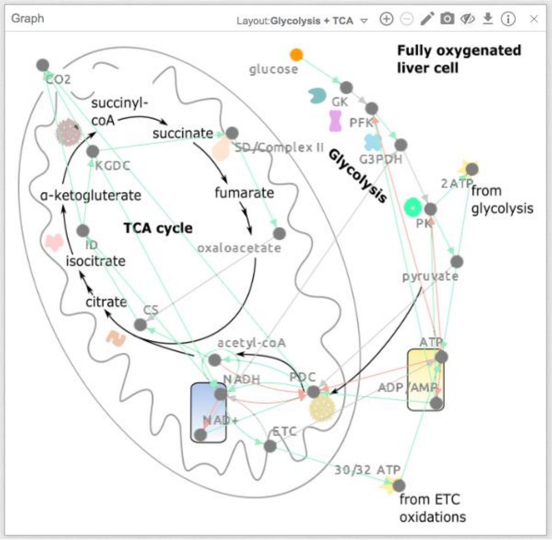


Next, students are presented with a model where the known feedback regulatory connections to TCA cycle enzymes are present (snapshot to the right - bottom). Through a series of questions, students determine which molecules negatively regulate which enzymes and then simulate the behavior of the updated model to compare the results to the previous simulation. Students evaluate the effect of the newly added connections on redox balance and on the production of glycolytic metabolites.

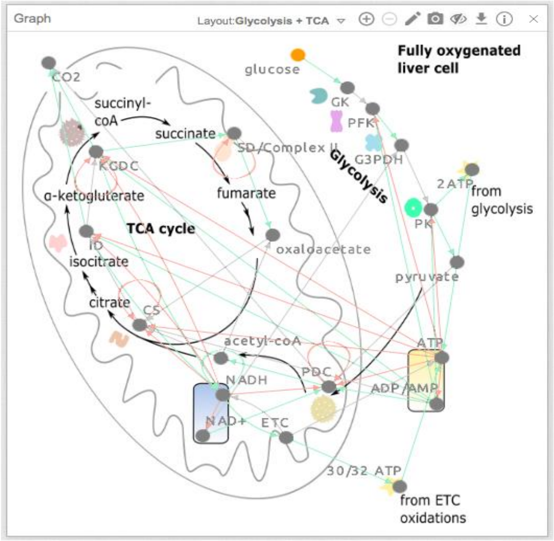


#### How instructors can help

Before students start the module:

1. Remind them to add the module to My Learning (clicked ADD TO MY LEARNING), via the Overview page of the module to enable the module to be edited.
2. Remind them to proceed via the Learning Activities page.

Remind students to pay attention to the cell type and oxygenation status of the cell represented by the model. Conceptual questions throughout the activities will require students to critically assess the purpose of homeostasis in the organism. Instructors can help by reminding them to draw from previous biology experience or discuss their thinking with a partner/instructor.

#### Model simulation review

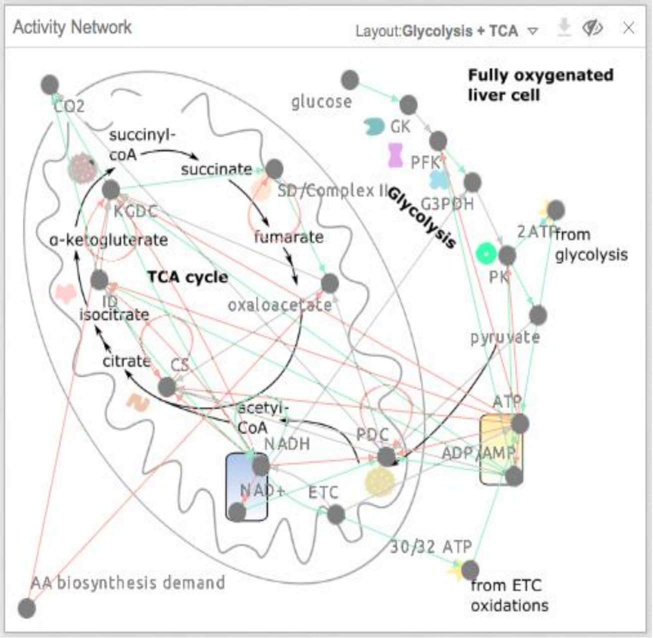


1. Energy charge (ADP and ATP) regulates TCA cycle enzymes.
2. NADH and metabolites regulate TCA cycle enzymes through product inhibition to maintain cellular homeostasis.
3. NAD+ and metabolites regulate TCA cycle enzymes through substrate availability to maintain cellular homeostasis.

The goal of the second half of the investigation is to introduce students to the **ability of an anaplerotic reaction to maintain the levels of TCA cycle metabolites**.

Students are presented with a computational model that allows the levels of amino acids being used by the cell to be manipulated externally. Students are asked to predict the simulation results from the model as amino acid demand changes from low to high and evaluate their results using simulation.

Next, students are asked to add a connection that represents an anaplerotic reaction and simulate the behavior of the model again as amino acid demand changes from low to high to demonstrate how anaplerotic reactions maintain TCA cycle metabolite levels.

#### How instructors can help

Although the questions are designed to focus student's attention on the model, it may be helpful to explicitly focus students' attention on the fact that the model changes as the scope of the questions being answered with the model expands.

#### Model connection building/simulation review (continued from the first half of the investigation)

4. Anaplerotic reactions can refill TCA cycle intermediates.
5. The TCA cycle can then keep running even when TCA cycle intermediates are needed for other cellular processes

#### Part 3: ETC and Fermentation

(Module ID: 29564 at https://cellcollective.org)

The diagram below shows the components of cellular respirations that are covered in this module:

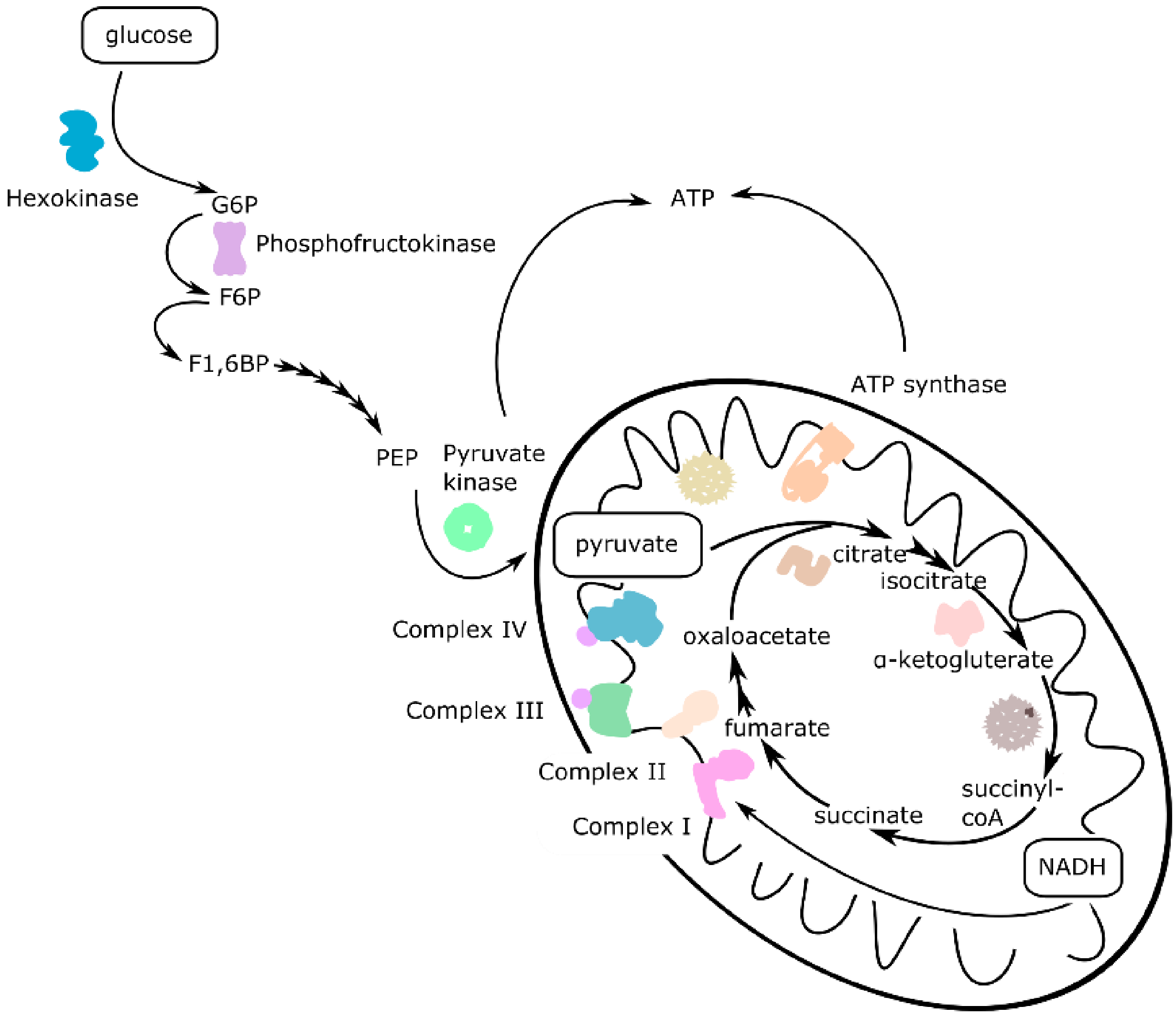

The goal of the investigation is to integrate concepts of energy charge- and redox-based regulation of glycolysis and the TCA cycle with electron transport chain function and cellular respiration.

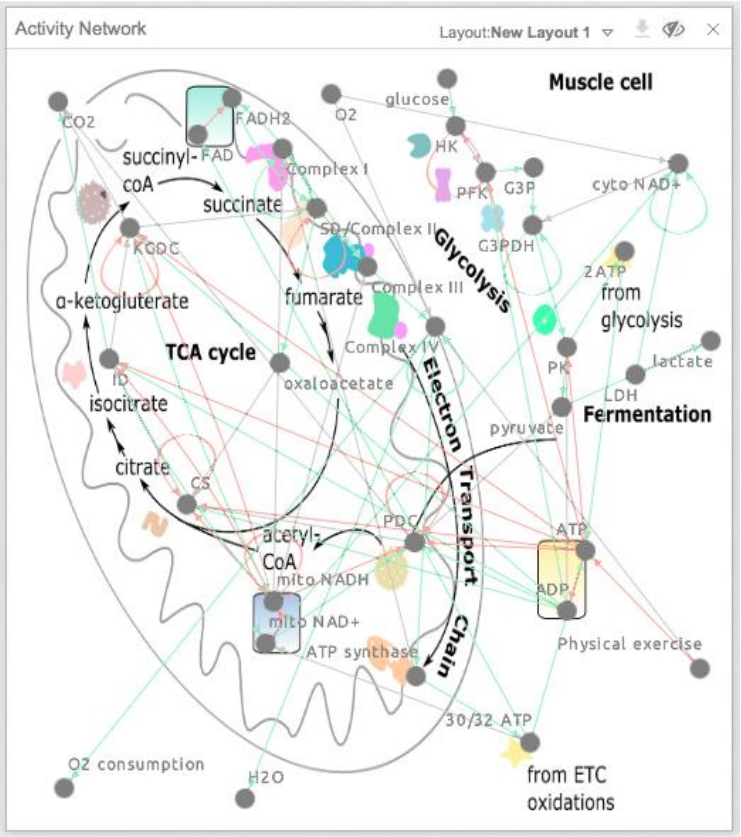


Students are presented with a computational model showing select enzymes and metabolites of glycolysis, the TCA cycle and the ETC with all known regulatory connections present (snapshot to the right).

Students are asked to predict the behavior of the model under three conditions: 1) oxygen present, no LDH expressed; 2) oxygen absent, no LDH expressed; and 3) oxygen absent, LDH expressed. Specifically, students are asked to predict the levels and activities of pyruvate and lactate, $CO_2$ production, $O_2$ consumption as readouts of the activity of specific cellular processes. Students also predict the levels of energy and redox molecules. Students are asked to record their predictions in tables. Next, they simulate the behavior of the model, tabulate the simulation results, and compare these results to their predictions. Finally, students are asked to critically evaluate the simulation results and explain how and why the system components and connections allow the cell to maintain homeostasis. Students then repeat a similar task, but this time they investigate the effect of exercise on the cell.

#### How instructors can help

Before students start the module:

1. Remind them to add the module to My Learning (clicked ADD TO MY LEARNING), via the Overview page of the module to enable the module to be edited.
2. Remind them to proceed via the Learning Activities page.

As before, students will be asked many conceptual questions throughout the activities that will require them to critically assess the purpose of homeostasis in the organism. It may be helpful to remind them to draw from previous biology experience or discuss their thinking with a partner/instructor.

#### Model simulation review

1. The cell can adjust its metabolism to oxygen availability and exercise through the coordinate regulation of glycolysis, the TCA cycle and ETC by enzymes “sensing” the levels of NADH/NAD+ and ATP/ADP.
2. When oxygen is limited, lactate dehydrogenase replenishes the cytoplasmic NAD+ pool so that glycolysis can proceed.
3. By allowing glycolysis to proceed when oxygen is absent (fermentation), the cell can produce some ATP. It is a lot less compared to oxidative respiration, so this is not a sustainable mode of ATP production for long periods of time.
4. When the cell begins to exercise and the ATP pool is constantly being depleted, it will increase glycolysis by relieving the inhibition on the rate-limiting enzymes of glycolysis. This allows the cell to increase glycolysis and oxidative respiration for ATP production.
5. After some time, oxygen will become depleted, but ATP production can be sustained for a short period of time by fermentation.

**File S4**

### Regulation of Cellular Respiration and Fermentation Assessments

#### Assessment 1.1: Glycolysis

1. Evaluate the following statements that describe the regulation of glycolytic enzymes (T/F):
   - A. T or F  Activation of pyruvate kinase by ADP maintains production of ATP.
   - B. T or F  Product inhibition of liver glucokinase would deregulate glucose storage.
   - C. T or F  Inhibition of muscle hexokinase by its product ensures that blood glucose is not wasted.
   - D. T or F  Glycolytic enzymes are regulated by energy charge to maximize energy production from glucose.
   - E. T or F  Liver glucokinase will be highly active under low blood glucose conditions.
   - F. T or F  Glycolytic flux in the liver cell is determined by the activity of glucokinase.
   - G. T or F  Glycolytic flux in the muscle cell is determined by the energy requirements of the cell.
   - H. T or F  Regulation of glycolysis by ADP increases the rate of glycolysis when energy is low.
   - I. T or F  Regulation of glycolysis by ADP and ATP stabilizes energy production when blood glucose varies.

#### Assessment 1.2: TCA

2. Evaluate the following statements that describe the regulation of the tricarboxylic acid (TCA) cycle enzymes (T/F):
   - A. T or F  Inhibition of pyruvate dehydrogenase complex decreases ATP production.
   - B. T or F  Regulation of TCA cycle enzymes allow anaplerotic reactions to refill the cycle.
   - C. T or F  TCA cycle enzymes are regulated by energy charge to maintain energy homeostasis.
   - D. T or F  NADH levels would remain unchanged if ATP began to accumulate.
   - E. T or F  Positive regulation of TCA cycle enzymes increases the levels of TCA metabolites.
   - F. T or F  Flux through the TCA cycle would decrease if NADH began to accumulate.
   - G. T or F  Anaplerotic reactions ensure that ATP production can proceed regardless of cellular amino acid demand.
   - H. T or F  Anaplerotic reactions ensure that cellular NADH levels are maintained.

#### Assessment 1.3: ETC and fermentation

3. Evaluate the following statements that compare respiration to fermentation and describe the regulation of the enzymes of the electron transport chain (ETC) (T/F):
   - A. T or F In the absence of $O_2$, glycolysis will be active if NAD+ levels can be maintained.
   - B. T or F The tricarboxylic acid (TCA) cycle will be active in the absence of $O_2$.
   - C. T or F ATP directly inhibits the enzymes of the electron transport chain.
   - D. T or F Activity of ETC enzymes would remain unchanged if ATP began to accumulate.
   - E. T or F Activity of ETC enzymes would increase if NADH began to accumulate.
   - ~~F. T or F If FADH2 increases, ETC activity decreases because of succinate dehydrogenase/complex II.~~
   - G. T or F Complex IV of the ETC will be active in the presence of $O_2$.
   - H. T or F NADH levels will increase in the absence of $O_2$.
   - I. T or F In the presence of $O_2$, ATP production is maintained by turning pyruvate into lactate.
   - J. T or F In the absence of $O_2$, less ATP production occurs.

**Note:** Please contact the authors to obtain a revised version showing wording updates and that corresponds to the latest online lesson. Item 3F had negative discrimination for both Biochem I courses and was not included in the analysis.

**File S5**

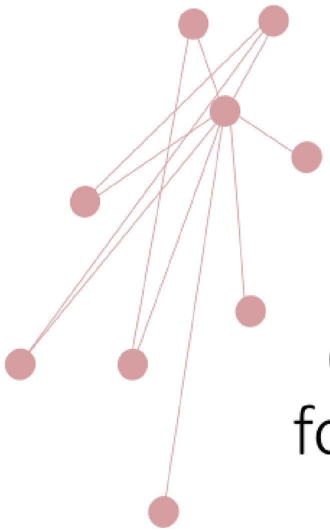

### A short introduction to computational modeling for biochemistry students

#### What is a ”system”?

“ a regularly interacting or interdependent group of items forming a unified whole”

- Merriam-Webster dictionary

There are many different kinds of systems:

- River system
- Digestive system
- Thermodynamic system
- Computer system

### How do we know that we understand the system?

By seeing how well we can **predict** that system's behavior:

If we can predict which medicine will relieve our digestive discomfort, we know we understand something about the digestive system and the causes of digestive discomfort.

### What about really complex biological systems, like cells?

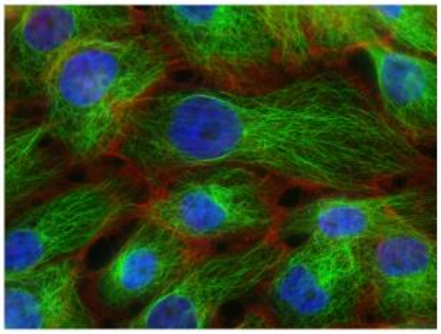

Photo credit: National Cancer Institute

We can reduce the system to smaller pieces, but keeping track of all the parallel processes is overwhelming

#### Even more than this…

Cells are not static, they are **dynamic**, and things are constantly changing.

This becomes even more challenging to try to predict how a cell will **dynamically** respond when something changes.

To predict how **many components** of a **complex system** change **dynamically** over time, one good solution is to:

1. Build a model of the biological system
2. Use a computer to simulate and/or analyze biological processes

#### When we have good predictive models of biological systems, we can…

- Understand how individual components interact together as a system
- Understand how the system responds to different conditions
- Understand which parts of the system are most critical for its function
- Identify drug targets and drug response

#### Purpose of the in-class modules:

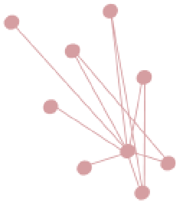

Using a computational modeling platform (Cell Collective Learn), you will explore important biochemical systems/processes and their regulation

#### In-class activity: Model purine biosynthesis regulation

https://learn.cellcollective.org/

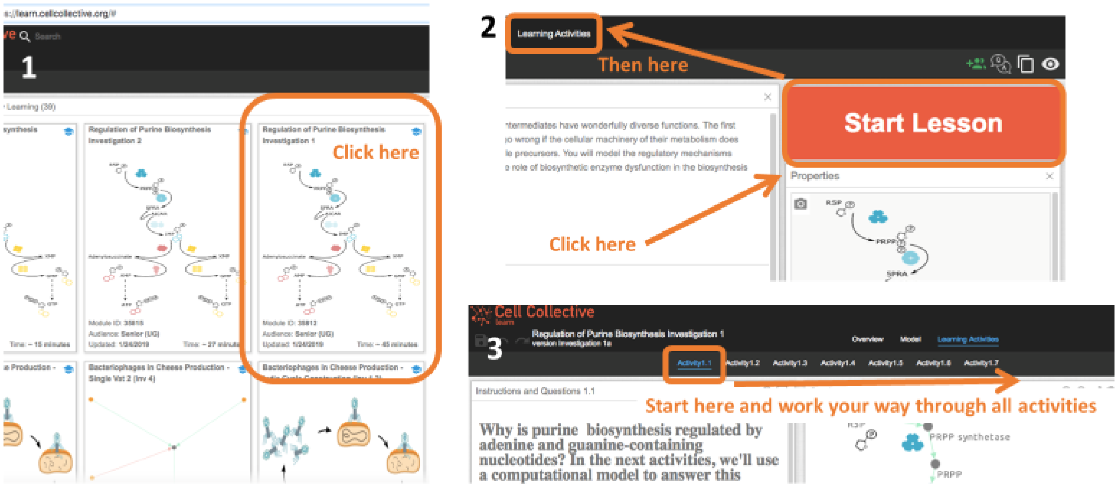

#### In-class activity: Model purine biosynthesis regulation

https://learn.cellcollective.org/

**What's different about arrows in the mathematical model compared to a typical biochemical diagram?**

**Using this kind of computational modeling approach:**

**A green arrow can represent ANY positive relationship (forward reaction, protein interaction, etc.)**
**A red arrow can represent ANY negative relationship (allosteric inhibition, reverse reaction, etc.)**

**Why are only some of the components modeled?**

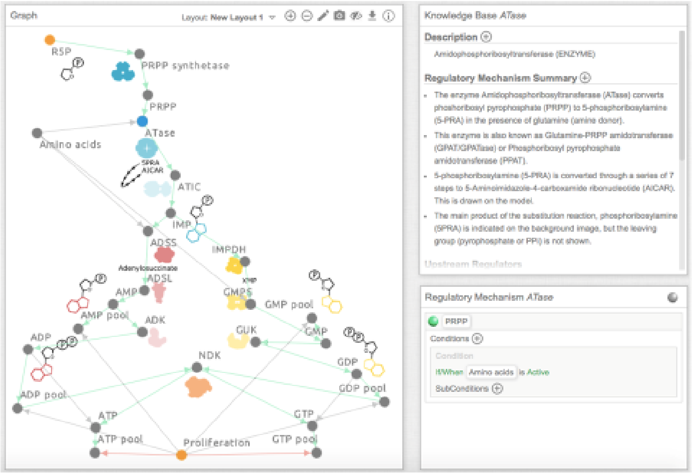


#### In-class activity: Model purine biosynthesis regulation

https://learn.cellcollective.org/

**<u>Some tips:</u>**
**You can only change dots (nodes) and arrows (edges), not images.**

**Details for each component is found in the Knowledge Base panel only if you click on the component – here you can find the details of how components are connected in the model.**

**If you want to remember the details or study for the exam, you can always come back to the first activity where the model AND the Knowledge base are.**

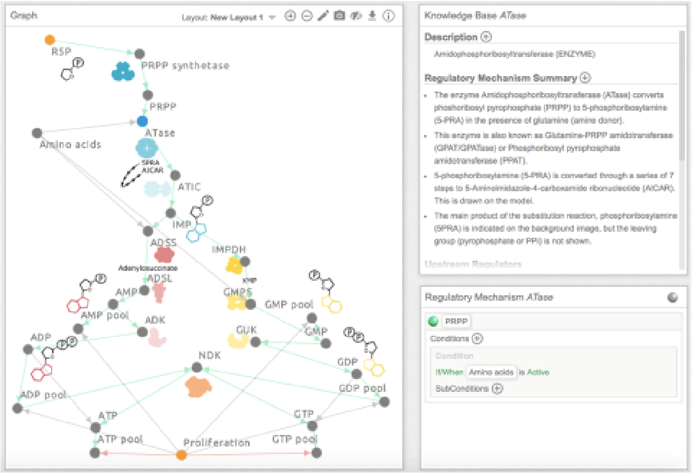

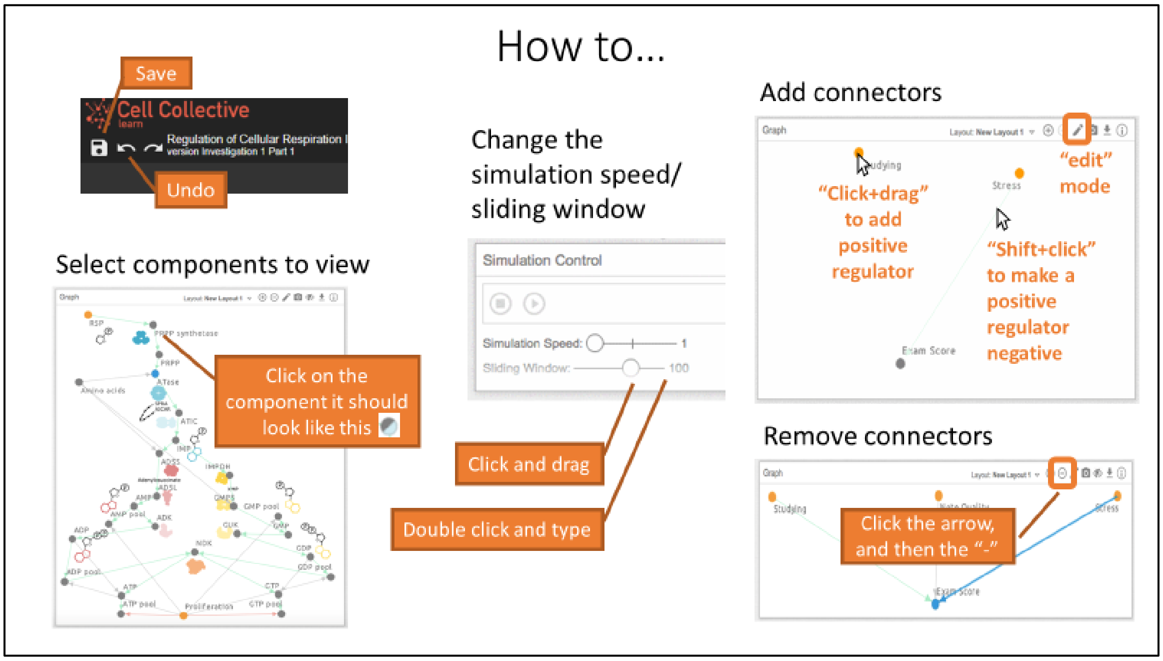
How to…
Save
Cell Collective
Regulation of Cellular Respiration
version Investigation 1 Part 1
Undo
Select components to view
Click on the component it should look like this
Change the simulation speed/ sliding window
Simulation Control
Simulation Speed:
Sliding Window:
100
Click and drag
Double click and type
Add connectors
Graph
"Click+drag" to add positive regulator
"edit" mode
Stress
"Shift+click" to make a positive regulator negative
Exam Score
Remove connectors
Studying
Click the arrow, and then the "-"

**File S6**

### Regulation of Purine Biosynthesis

### Instructor Guide

#### Part 1: Negative Allosteric Feedback

(Module ID: 35812 at https://cellcollective.org)

The diagram below shows the components of purine biosynthesis that are covered in this module:

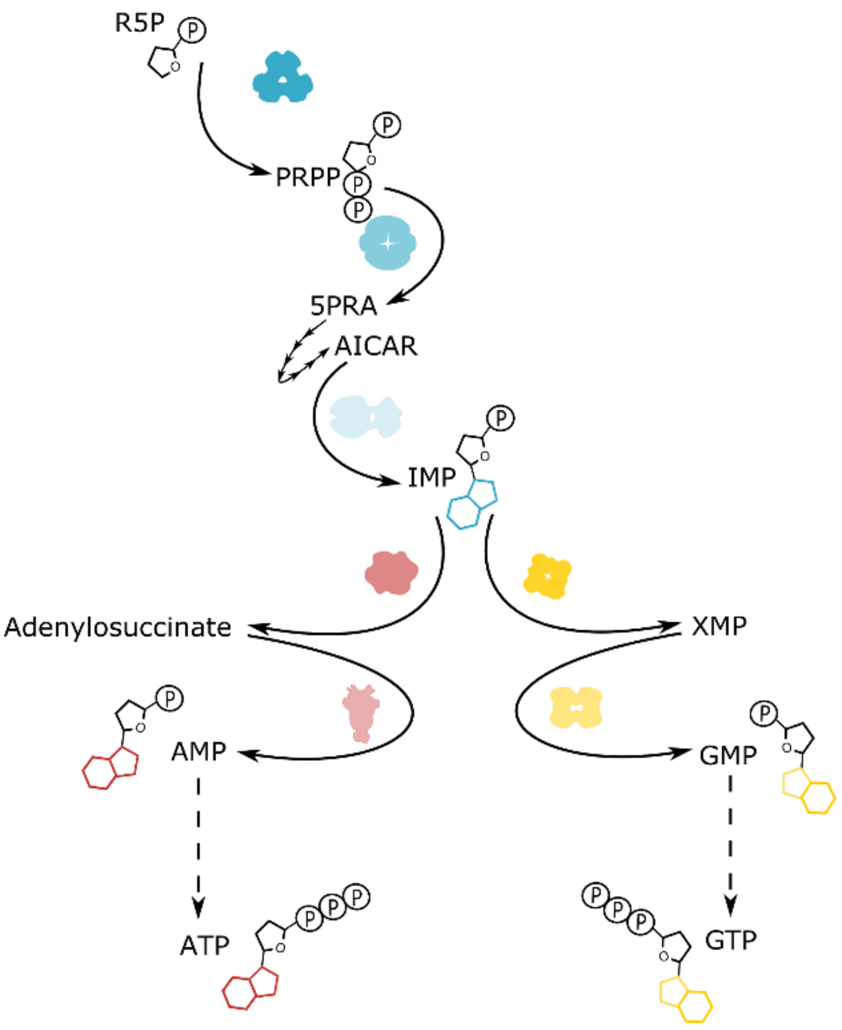

The goal of the first half of the investigation is to introduce students to the importance of **nucleotide-based regulation of purine biosynthetic enzymes** to maintain **nucleotide homeostasis**.

Students are presented with a computational model showing select enzymes and metabolites of purine biosynthesis and most of the known allosteric feedback regulatory connections are not present (snapshot to the right).

Students are asked to simulate the behavior of the model as is (snapshot to the right above) and then to add negative allosteric feedback relationships from the ADP and GDP pools to PRPP synthetase, which catalyzes a rate-determining step of purine biosynthesis. They simulate the behavior of the model after adding these regulatory connections and tabulate the simulation results. Students are then provided with a model showing all the known allosteric regulatory connections (snapshot to the far right above). Again, they simulate the model behavior and tabulate the results. Finally, they evaluate their various simulation results. Throughout the investigation, students are asked to reason about how these regulatory connections will affect the entire organism.

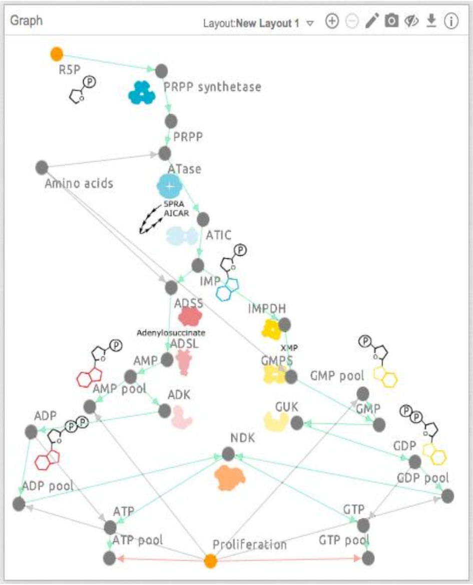


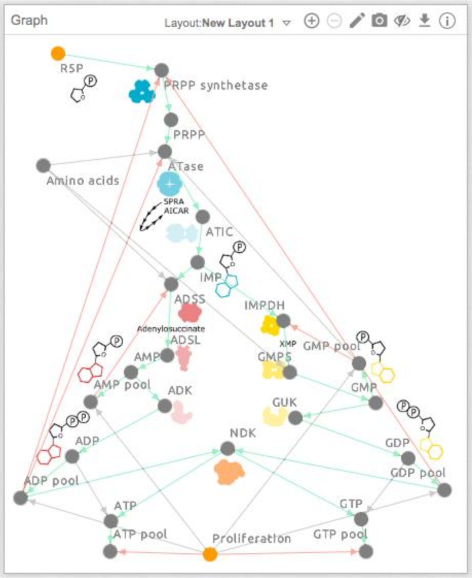


#### How instructors can help

Before students start the module:

1. Remind them to add the module to My Learning (click the “Start Lesson” button), via the Overview page of the module. This will enable the module to be edited.
2. Direct them to the Learning Activities page in Cell Collective.
3. Ask students to confirm that the model is in “edit” mode (this should be the default and is indicated by a “pencil” icon, however, if the model is in “view” mode, students can click the “eye” icon within the Graph panel to change it to “edit” mode).
4. Remind students to:
   a. To draw a connection (arrow), click the starting component, drag, and release the mouse over the component you want the arrowhead to land on.
   b. To delete an arrow, highlight it, then press delete (fn+delete for mac).
   c. To toggle an arrow from positive to negative: (1) deselect everything! (2) press and hold shift while you next click on the arrow that you want to change. Click somewhere else to see the effect.
5. If students need to return to their lesson, remind them to access it through “My learning”, not “Public modules”.

Although this is pointed out explicitly in the lesson, it may still be important to remind students the model focuses on rapid control mechanisms only and that other cellular processes, such as respiration, although not explicitly modeled, should not be completely ignored. It may be important to continually remind students that models frequently present incomplete views of a complex reality. Students will be asked many conceptual questions throughout the activities that will require them to critically assess the purpose of homeostasis in the organism. In general, it may be helpful to remind them to draw from previous biology experience or discuss their thinking with a partner/instructor.

##### Model connection building/simulation review

1. ADP and GDP negatively regulate purine biosynthetic enzymes (PRPP synthetase, ATase, ADSS, and IMPDH) which reduces the levels of IMP through the biosynthesis pathway.
2. Nucleotide-based regulation by ADP and GDP ensures that the cell has sufficient nucleotide supply and can respond to increased demand for nucleotides (maintains homeostasis!) regardless of how much R5P is available.

#### Part 2: "Cross-regulation" of purine branches

(Module ID: 35815 at https://cellcollective.org)

The same components of purine biosynthesis as Part 1 are covered in this module (diagram on first page of guide).

The goal of this investigation is to introduce students to the importance of **"cross-regulation" of the two branches of purine biosynthesis** and **how substrate availability from one branch is able to balance nucleotide levels in the other branch**.

Students are presented with a table of simulation results that were previously obtained using versions of the model where more regulatory interactions were sequentially added to the system and the simulation results were recorded. This first activity is an extension and review of Part 1 of the lesson, and students are asked to use similar reasoning to explain the results as they used for Part 1.

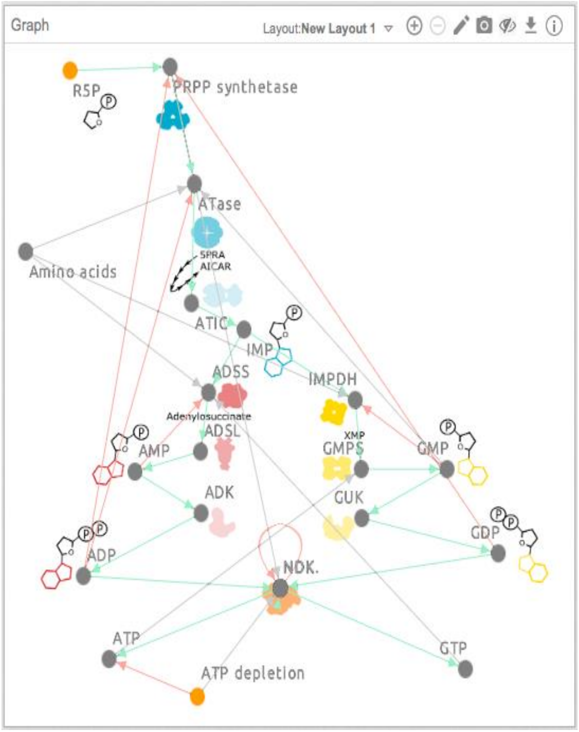


Next, students are presented with a model where the "cross-regulatory" interactions are added (snapshot to the right). They are asked to predict what would happen to ATP and GTP levels when adenine-rich DNA must be made. They simulate the model, test their predictions, and reason through why the result is the way it is. To be successful, they must rely on their previous knowledge of how substrate availability affects the enzymatic rate.

#### How instructors can help

Before students start the module:

1. Remind them to add the module to My Learning (click the “Start Lesson” button), via the Overview page of the module to enable the module to be edited.
2. Direct them to the Learning Activities page in Cell Collective.
3. If students need to return to their lesson, remind them to access it through “My learning”, not “Public modules”.

Remind students to recall their results and the concepts they covered in Part 1. Conceptual questions throughout the activities will require students to critically assess the purpose of homeostasis in the organism. Instructors can help by reminding them to draw from previous biology experience or discuss their thinking with a partner/instructor.

#### Model simulation review

1. Purines regulate multiple enzymes that determine the production of intermediates that are required for their synthesis.
2. The regulation occurs at various positions in the pathway to ensure redundancy.
3. Biosynthesis of adenine and guanine nucleotides is interrelated to ensure that the levels of both types of nucleotides remain balanced within the cell.

### Part 3: Negative Allosteric Feedback

(Module ID: 35819 at https://cellcollective.org)

The same components of purine biosynthesis as Part 1 are covered in this module (diagram on page 1).

The goal of the investigation is to conceptually integrate the process of purine *de novo* biosynthesis within the larger context of the cell to include other important purine-related processes such as purine degradation and salvage. These ideas are further extended by asking students to evaluate the effect of different mutations of *de novo* biosynthetic enzymes on cellular purine levels, the cell, and the organism.

Students are presented with diagrams (snapshots to the right) that conceptually extend the concepts they have already learned using the computational models in Parts 1 and 2 of the lesson.

Students simulate the model and evaluate the simulation results when 1) all the enzymes of purine biosynthesis are wild-type enzymes and when there are mutations in two key enzymes of *de novo* biosynthesis: 2) an activating mutation in PRPP synthetase; and 3) an inactivating mutation in adenylosuccinate lyase (ADSL). Students are asked to record their results in tables and critically evaluate the simulation results and explain how the cell could compensate when mutant enzymes are expressed.

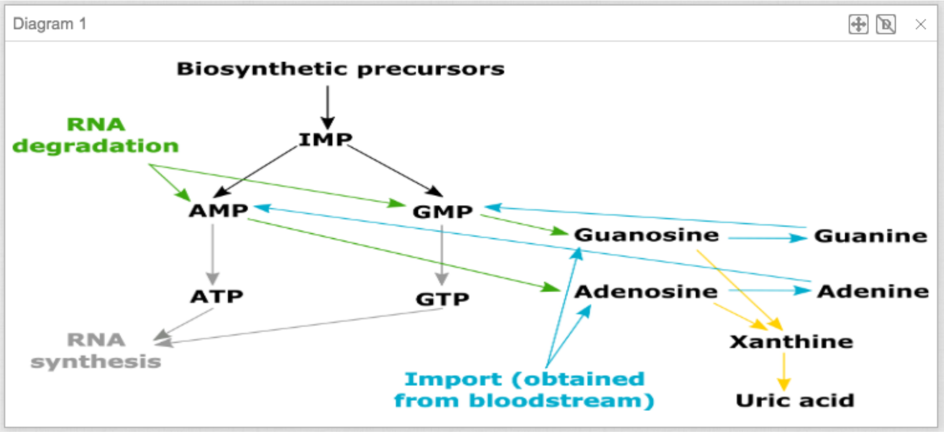


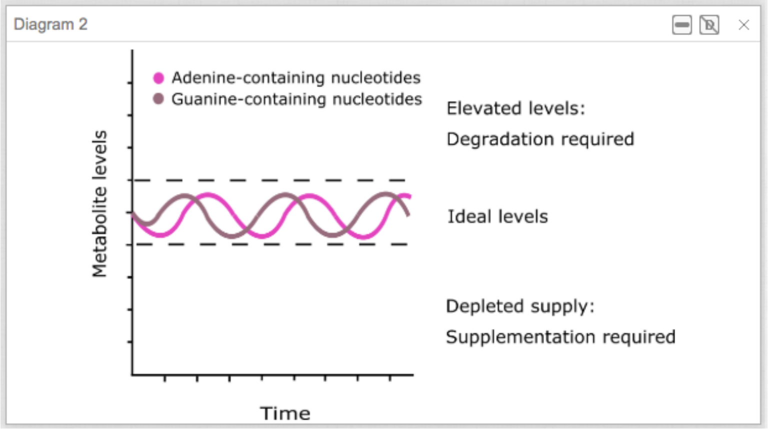


#### How instructors can help

Before students start the module:

1. Remind them to add the module to My Learning (click the “Start Lesson” button), via the Overview page of the module to enable the module to be edited.
2. Direct them to the Learning Activities page in Cell Collective.
3. If students need to return to their lesson, remind them to access it through “My learning”, not “Public modules”.

As before, students will be asked many conceptual questions throughout the activities that will require them to critically assess the purpose of homeostasis in the organism. It may be helpful to continue reminding them to draw from previous biology experience or discuss their thinking with a partner/instructor.

#### Model simulation review

1. Mutations in purine biosynthetic enzymes adversely affect cellular purine levels and these effects must be compensated by changing the activity of other cellular processes such as purine salvage and degradation.
2. Specifically, PRPP synthetase overactivity can override feedback regulation, causing accumulation of purine nucleotides that must be degraded. Conversely, ADSL deficiency reduces nucleotides in the cell, and dietary supplementation will be required.

**File S7**

### Regulation of Purine Biosynthesis

### Assessment 2.1: Purine Biosynthesis

1. Evaluate the following statements describing interactions between the elements of *de novo* purine biosynthesis (T/F):
    - ~~A. T or F Two enzymes in the main branch of *de novo* biosynthesis are feedback inhibited~~
    - B. T or F Two enzymes in the GTP branch of *de novo* purine biosynthesis are feedback inhibited.
    - C. T or F IMP dehydrogenase (IMPDH) is regulated by allosteric feedback inhibition.
    - D. T or F GMP synthetase is regulated by allosteric feedback inhibition.
    - ~~E. T or F Glutamine PRPP amidotransferase (ATase) is regulated by substrate availability.~~
    - F. T or F IMP is a precursor only for GTP biosynthesis.
    - G. T or F Glutamine PRPP amidotransferase (ATase) is common to both ATP and GTP biosynthesis.
    - H. T or F Adenylosuccinate synthetase (ADSL) is common to both ATP and GTP biosynthesis.
    - I. T or F The levels of ATP and GTP in the cell determine the rate of GTP biosynthesis.

2. Determine whether the following statements describe how the regulation of de novo purine biosynthesis is integrated to maintain homeostasis (T/F):
    - A. T or F ATP can only be produced through de novo biosynthesis when both PRPP and GTP are present.
    - B. T or F If PRPP synthetase is not regulated by ADP and GDP, PRPP could accumulate in the cell and potentially become toxic.
    - C. T or F If IMP dehydrogenase (IMPDH) is not regulated by GMP, both ATP and GTP would accumulate in the cell and potentially become toxic.

3. In an actively proliferating cell, the following describe the *de novo* purine biosynthesis pathway (T/F):
    - A. T or F ATP and GTP will be overproduced to meet cellular demands, and remain high.
    - B. T or F ATP and GTP levels will initially fall, but return to normal as the allosteric inhibition on biosynthetic enzymes is relieved.
    - C. T or F Metabolic flux through de novo purine biosynthesis will temporarily increase to accommodate cellular demand for ATP and GTP synthesis.
    - D. T or F ATP and GTP levels will initially fall, and will remain low while the cell is proliferating.

4. The following statements describe the effect of mutations of the enzymes of *de novo* purine biosynthesis (T/F):
   For activating mutations in PRPP synthetase, the following may be expected:
   A. T or F Increased production of nucleotides.
   B. T or F Increased flux through salvage pathways to compensate for metabolic imbalances.
   C. T or F Compensatory pathways could somewhat mitigate the effects on nucleotide levels.

   For inactivating mutations in Adenylosuccinate lyase (ADSL), the following may be expected:
   A. T or F Decreased production of nucleotides.
   B. T or F Increased flux through degradation pathways to compensate for metabolic imbalances.
   C. T or F Compensatory pathways could completely mitigate effects on nucleotide pathways.

**Note:** Items 1A and 1E had negative discrimination for the Biochemistry II course and was not included in the analysis.

**File S8**

### Regulation of Cellular Respiration and Fermentation

### Survey: student experiences with the lesson

Q1. Please comment on your learning after completing this module.

| | Strongly disagree | Disagree | Neither agree nor disagree | Agree | Strongly agree |
|---|---|---|---|---|---|
| a. The module helped me to understand how the regulation of glycolysis, the TCA cycle, and the ETC are integrated (how it works together) | ○ | ○ | ○ | ○ | ○ |
| b. The simulations were helpful to understand the effects of feedback loops and environmental conditions on the entire system | ○ | ○ | ○ | ○ | ○ |
| c. The simulations were helpful to understand the effects of feedback loops and environmental conditions on the entire system | ○ | ○ | ○ | ○ | ○ |
| d. The module helped me to remember to think about both the individual components and also their connection to the larger process | ○ | ○ | ○ | ○ | ○ |
| e. I think I will remember what I learned about the regulation of cellular respiration better than I would have if I did not complete the module. | ○ | ○ | ○ | ○ | ○ |
| f. I think I understand what I learned about the regulation of cellular respiration better than I would have if I did not complete the module. | ○ | ○ | ○ | ○ | ○ |
| g. Overall, completing this module assisted my learning of the material | ○ | ○ | ○ | ○ | ○ |

Q2. Please comment on which parts of the module you found most effective to aid your learning (which parts helped you the most).

Q3. Please comment on which parts of the module you found least effective to aid your learning (which parts helped you the least).

Q4. Please list one concept or idea that you are still unsure about after completing this module.

Q5. What was most challenging about working with the computational modules?

Q6. Knowing that you will still be responsible for understanding the regulation of cellular respiration, and that computational skills are important to develop for various reasons, how could the activity be changed to aid your learning?

Q7. Do you have any other feedback that you would like to provide?

**File S9**

### Regulation of Purine Biosynthesis

### Survey: student experiences with the lesson

Q1. Please comment on your learning after completing this module.

| | Strongly disagree | Disagree | Neither agree nor disagree | Agree | Strongly agree |
|---|---|---|---|---|---|
| a. The module helped me to understand how the regulation of purine biosynthesis maintains purine homeostasis regardless of changing cellular conditions | ○ | ○ | ○ | ○ | ○ |
| b. The simulations were helpful to understand the effects of specific feedback loops (the results of allosteric regulation) on ATP and GTP production | ○ | ○ | ○ | ○ | ○ |
| c. The module helped me to remember to think about both the individual components and also their connection to the larger process | ○ | ○ | ○ | ○ | ○ |
| d. I think I learned about the topic of regulation of purine biosynthesis in much greater depth than I would have if I did not complete the module | ○ | ○ | ○ | ○ | ○ |
| e. I think I understand what I learned about regulation of purine biosynthesis better than I would have if I did not complete the module.e if I did not complete the module. | ○ | ○ | ○ | ○ | ○ |
| f. Overall, completing this module assisted my learning of the material | ○ | ○ | ○ | ○ | ○ |

Q2. Please comment on which parts of the module you found most effective to aid your learning (which parts helped you the most).

Q3. lease comment on which parts of the module you found least effective to aid your learning (which parts helped you the least):

Q4.Please list one concept or idea that you are still unsure about after completing the module.

Q5. What was most challenging about working with the computational modules?

Q6. Knowing that you will still be responsible for understanding the regulation of purine biosynthesis, and that computational skills are important to develop for various reasons, how could the activity be changed to aid your learning?

Q7. Do you have any other feedback that you would like to provide?
